# Designed Peptides as Affinity Ligands for Extracellular-Vesicle-based Cancer Biomarker Detection

**DOI:** 10.1101/2025.03.27.645572

**Authors:** Sudeep Sarma, Young Kwan Cho, Arpan Tapdiya, Jin-Ho Park, Hakho Lee, Carol K. Hall

**Affiliations:** Department of Chemical and Biomolecular Engineering, North Carolina State University, Raleigh, NC, USA; Center for Systems Biology, Massachusetts General Hospital, Boston, MA 02114, USA; Department of Radiology, Massachusetts General Hospital, Harvard Medical School, Boston, MA, USA

**Keywords:** Peptide-based bioassay, Cancer detection, EV imaging, Tetraspanins, EpCAM

## Abstract

Assays for cancer diagnosis via the analysis of tumor biomarkers on circulating extracellular vesicles (EVs) have shown great potential. Single EV imaging that can measure the abundance of protein biomarkers in EVs can help in detecting the presence, stage, and progression of disease. Antibodies are typically used to detect EV proteins. Controlling the quality of antibody-based immunoassays can, however, be challenging, as they may exhibit unintended cross-reactivity with non-target proteins or variability in binding affinity across different batches, even for monoclonal antibodies. Here, we report short peptides that are a promising alternative to antibodies for detecting protein biomarkers in EVs. We describe an effort that combines a *Pep*tide *B*inding *D*esign (PepBD) algorithm and molecular-level simulations to identify peptides that can recognize (1) the extracellular domain of EpCAM (a known cancer biomarker) and (2) the extracellular domain of tetraspanin CD81(a protein commonly expressed on the surface of EVs). The peptides designed for EpCAM and CD81 were labeled with a fluorescent dye and their binding to the target protein was evaluated using fluorescence ELISA. The results demonstrated that one of the computationally designed peptides EP-2.1 exhibited an affinity for the target EpCAM protein comparable to that of an antibody. Further testing involved single EV imaging to gauge the peptide’s affinity towards EVs. Designed peptides for both the target proteins showed similar affinity as antibodies to EVs.

## 1. INTRODUCTION

Cancer remains a global health crisis, accounting for one in six deaths worldwide^1^. The disease is categorized into stages (I-IV) based on its spread from the primary infected organ, with approximately 50% of cases diagnosed at advanced stages (II-IV)^2,3^. Treatment options for advanced cancer are often limited, resulting in significantly lower survival rates. While stage I cancers have a five-year survival rate of 70–90%, this drops to 10–20% for stage IV cases^3^. Clearly, the identification of cancerous tumors at an early stage is important.

For early detection, an understanding of cancer progression is essential. Biomarkers are measurable indicators of different biological processes in the body^4^. They can help predict the onset of a disease, its progression and the likely outcome of treatment^5^. Protein biomarkers have proven to be pivotal in cancer diagnostics^6,7^. EpCAM, which is overexpressed in epithelial cell tumors, is found in 85% of adenocarcinomas and 72% of squamous cell carcinomas^8,9^. In ovarian cancer, EpCAM is overexpressed in up to 75% of epithelial ovarian carcinomas, particularly in aggressive subtypes^10,11^. Methods like liquid biopsies and immunoassays are used to find these biomarkers and hence enhance early detection of cancer.

Liquid biopsies are lab tests done on a sample of body fluids to look for circulating tumor cells, tumor DNAs, and extracellular vesicles, which may originate from cancerous cells^12^. Extracellular vesicles (EVs) are heterogeneous, membrane-bound phospholipid vesicles that are secreted by a variety of mammalian cells, including host cells and dividing cancer cells^13^. Although initially considered as “cell dust”, and a means to dispose of cellular components, they are now recognized as a circulating source of biomarkers from different diseases. EVs carry molecular cargo derived from tumor cells and are found circulating in readily accessible body fluids^13^. Tumor-associated EVs have been used effectively as biomarkers to define tumor type and malignancy stage (“liquid biopsies”). Immunoassays, another key diagnostic tool, rely on the interaction between antibodies and antigens to detect biomarkers. These assays are simple, rapid, and cost-effective, and can simultaneously detect multiple biomarkers^14^. However, controlling the quality of immunoassays based on antibodies has some challenges. Due to their large size, antibodies may exhibit accidental cross-reactivity with non-targeted proteins, resulting in background noise in the analytical signals^15^. Variability in the binding affinity of antibodies with the targeted protein is also observed^16^, potentially leading to inconsistent and non-reproducible results.

Short peptides, consisting of 8–15 amino acid residues, serve as a promising alternative to antibodies in immunoassays. Their small size minimizes cross-reactivity, enhances binding consistency, and enables dense surface immobilization, amplifying the signal output being measured during immunoassays. Additionally, peptides are cost-effective, easy to synthesize, and exhibit enhanced environmental stability compared to antibodies due to their simpler structures, thus making them less susceptible to degradation, and hence ideal for diagnostic applications^17,18^. *In-vitro* methods like phage display library screening^19^ and ELISA^14^ have been successful in identifying high-affinity peptides, but face drawbacks such as limited peptide library diversity due to constraints on library size and sequence complexity^20^. Phage display libraries may also have a selection bias towards those peptide sequences that are most efficiently displayed on the phage surface^21^. To address these challenges, computational peptide design has emerged as a method that enables the rapid exploration of the vast peptide sequence space to identify peptides with high specificity and affinity for target biomolecules (proteins, RNA etc.)^22^. These *in-silico* techniques can complement experimental approaches like phage display by guiding the selection of peptide candidates for further validation^23^. Integrating computational and experimental methodologies holds promise for advancing cancer diagnostics, enabling earlier detection and hence more effective treatment.

In this paper, we present a combined computational and experimental effort to discover synthetic peptide ligands that can detect cancer biomarkers. Specifically, we employ the Hall group’s computational *Pep*tide *B*inding *D*esign (PepBD) algorithm^24–27^ to design peptides and validate these peptides through an immunoassay-based EV-imaging technique developed by the Lee group. The discovered peptide ligands are intended to serve as biological recognition elements for biosensor devices, enabling early cancer detection. Our primary focus is on designing peptide ligands that recognize EpCAM (epithelial cell adhesion molecule, CD326), a well-established biomarker for ovarian cancer diagnosis^28^, and the tetraspanin protein CD81, a key biomarker for extracellular vesicle (EV) identification^13^. EpCAM is a conserved glycoprotein found on tumor-associated epithelial cells and EVs, while CD81 is a transmembrane protein abundantly expressed on EV surfaces, facilitating EV quantification and characterization. Our focus on ovarian cancer (OvCa) is driven by the high percentage of OvCa cases being diagnosed at advanced stages, and the high capacity of OvCa tumors to metastasize, making ovarian cancer the most lethal gynecological cancer^29^.

To achieve our goal of discovering peptide ligands for biomarkers associated with OvCa, we first computationally design peptide ligands using PepBD. PepBD, a peptide discovery algorithm, has been successfully applied to design peptides that bind to various target proteins, including transfer RNALys3^24^, cardiac biomarkers like troponin I^30^ and neuropeptide Y^31^, immunoglobulin G domains^32^, the SARS-CoV-2 spike protein^33^, and toxins in Clostridioides difficile infection^34^. The PepBD algorithm is based on applying the Monte-Carlo method to explore the vast conformational and sequence space of peptides. The algorithm requires a starting input complex between a known peptide ligand (referred to as the reference peptide) and the target protein, which in this study are EpCAM and CD81. A reference peptide is a previously-identified sequence known for its ability to bind to a particular target protein. A score function, 𝛤_𝑠𝑐𝑜𝑟𝑒_, that measures the binding energy of the peptide to the receptor and the conformational stability of the peptide when bound to the target protein is used to accept or reject newly-generated peptide sequences.

The PepBD algorithm allows us to design variants of the reference peptides that can bind with high affinity to the target proteins EpCAM and CD81. By employing two distinct peptide probes, one targeting tetraspanins for EV identification and the other targeting EpCAM, we aimed to develop a robust method for detection of biomarkers associated with OvCa. For this study, the starting reference peptides for the protein EpCAM include two 9-mer sequences, EP-1 (YEVHTYYLD) and EP-2 (VSVHTYDLE), which were identified by Weizhi et al^35^. The starting reference peptide for the protein CD81 is P152 (CFMKRLRK), an 8-mer peptide identified by Suwatthanarak et al^36^. Weizhi et al. and Suwatthanarak et al. both used high-throughput microarray-based peptide screening methods to identify their peptides^35,37^. We first performed docking simulations between the reference peptides and the target proteins to generate initial structures for seeding PepBD designs. The selection of the best docked complexes was based on their binding free energies, which were calculated using explicit-solvent molecular dynamics (MD) simulations in the AMBER20 package^38^ and the MM/GBSA protocol^39^. Several PepBD runs were conducted to design new peptide sequences, which were further evaluated using similar explicit-solvent MD simulations and free energy calculations. To experimentally characterize the binding capabilities of the computationally designed peptide ligands towards the target proteins, we utilized immunoassays and single-EV fluorescent imaging .

## 2. RESULTS

### 2.1 Computational design of peptides for EpCAM

#### 2.1.1 Structure of the reference peptides bound to the target protein EpCAM

We began by determining the structures of peptides EP-1 (sequence: YEVHTYYLD) and EP-2 (sequence: VSVHTYDLE), which were identified as being able to bind to EpCAM by Weizhi et al^35^. The complexes that form when these structures are bound to EpCAM can serve as starting inputs for our design process. The crystal structure of EpCAM corresponding to PDB ID: 4MZV^40^ is shown in **Figure 1A**. We refined this structure by performing a brief 5 ns MD simulation. Two putative binding regions on EpCAM were identified: (1) the N terminal domain – NTD (shown as red in **Figure 1A**) and (2) the pocket P1 (shown as green in **Figure 1A**). The NTD on EpCAM was chosen as a binding site as it is targeted by the vast majority of anti-EpCAM monoclonal antibodies^41^. Pocket P1 was also selected as an additional binding site as it was predicted to be the most likely binding pocket from an *in-silico* druggability analysis that we performed on the crystal structure of EpCAM using PockDrug^42^. The coordinate files for peptides EP-1 and EP-2 were generated by performing molecular dynamics simulations in the AMBER20 suite using the ff14SB forcefield^43^. Briefly, a 5ns MD simulation of the peptide in a simulation box with periodic boundary conditions containing ∼3200 water molecules was conducted. Peptides EP-1 and EP-2 were docked *in silico* against the NTD and against the pocket P1 on the crystal structure of EpCAM using the docking software HADDOCKv2.4^44,45^. The docking simulations generated by applying HADDOCKv2.4 on the four peptide-EpCAM complexes, EP-1-EpCAM NTD, EP-1-EpCAM P1, EP-2-EpCAM NTD and EP-2-EpCAM P1 produced multiple binding poses. These poses were clustered based on the fraction of common contacts, where a “cluster” was defined as a collection of at-least four different structures with > 80% similar contacts. For each of the four peptide- EpCAM complexes, the binding pose from the first cluster was identified as the best binding conformation and thus, was selected for further refinement and evaluation using MD simulations. Three independent explicit-solvent 100 ns atomistic molecular dynamics simulations were carried out for the four complexes: EP-1-EpCAM NTD, EP-1-EpCAM P1, EP-2-EpCAM NTD and EP- 2-EpCAM P1. After the MD simulations, the Δ𝐺_binding_of the peptide-receptor complex was calculated using the MM/GBSA protocol^39^ and variable dielectric constant methods^46^. Details of our MD simulations and Δ𝐺_binding_calculation procedures are provided in *Methods and Materials (*>*section 4.2)*.

**Figure 1.**
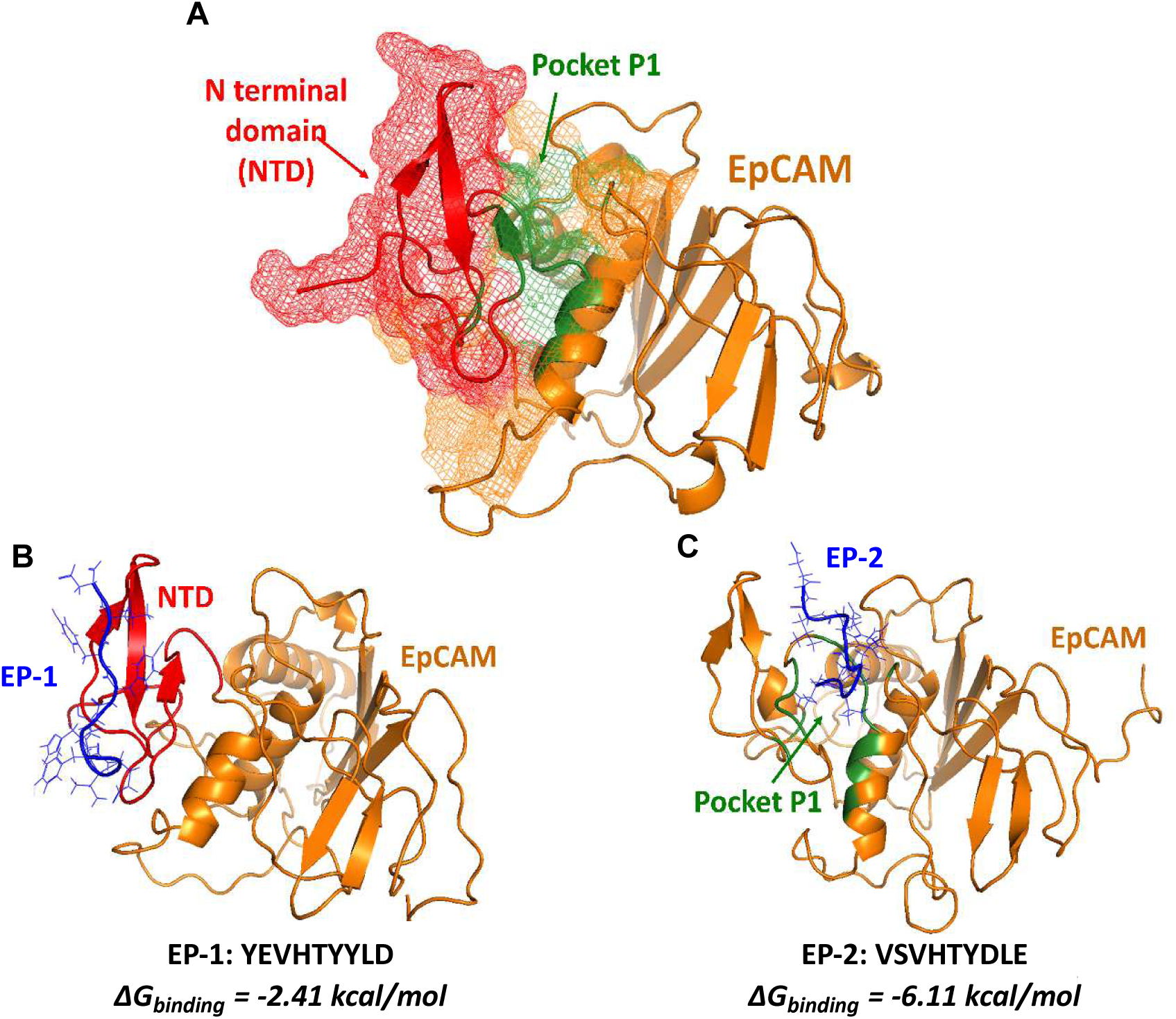
(A) Th**e** crystal structure of EpCAM (orange) with the N-terminal domain (NTD, red) and the putative binding pocket P1 (green) identified by using the PockDrug server (PDB ID: 4MZV). **(B)** Peptide EP-1 (sequence: YEVHTYYLD, blue) bound to the NTD of the EpCAM protein and **(C)** peptide EP-2 (sequence: VSVHTYDLE, blue) bound to pocket P1 of the EpCAM protein.

The simulation results suggested that EP-1 binds to the NTD with Δ𝐺_binding_ = -2.41 kcal/mol) and to the pocket P1 with Δ𝐺_binding_= -2.13 kcal/mol) on EpCAM, while EP-2 only binds to the pocket P1 with Δ𝐺_binding_ = -6.11 kcal/mol) on EpCAM. Table 1 summarizes the results from the molecular dynamics simulations of EP-1 and EP-2 interacting with EpCAM. Since EP-1 had a higher binding affinity for NTD than for pocket P1, while EP-2 only binds to pocket P1, the EP-1-EpCAM NTD complex (**Figure 1B**) and EP-2-EpCAM P1 complex (**Figure 1C**) were selected as input structures for PepBD optimization.

**Table 1.** The Δ𝐺_binding_values of the peptides EP-1 and EP-2 binding to the NTD and to the pocket P1 on EpCAM obtained from MD simulations.

| *Peptide-protein complex* | *$\Delta G_{binding}$ (kcal/mol)* |
| --- | --- |
| EP-1-EpCAM NTD | -2.41 |
| EP-1-EpCAM P1 | -2.13 |
| EP-2-EpCAM NTD | Does not bind ($\Delta G > 0$) |
| EP-2-EpCAM P1 | -6.11 |

#### 2.1.2 Design of EpCAM-binding peptides using PepBD and evaluation of binding free energies

Based on the results of our MD simulations, we first determined which fragment on peptides EP-1 and EP-2 play an important role in binding to EpCAM. Specifically, for EP-1 binding to the NTD region of EpCAM, the C-terminal residues contributed significantly more to the interaction energy than the flanking N-terminal residues (as indicated in **Supplementary Figure S1A**). Similarly, for EP-2 binding to pocket P1 on EpCAM, the N-terminal residues had a higher contribution to the interaction energy than the flanking C-terminal residues (as indicated in **Supplementary Figure S1B**). We added -GSG- residues (a non-reactive segment) to the N-terminus of EP-1 and to the C-terminus of EP-2 and performed independent 100 ns MD simulations of the GSG-EP-1-EpCAM NTD complex and of the EP-2-GSG-EpCAM P1 complex. The non-reactive segment was attached to the peptides in the computational design to mimic the spacer molecule that couples the peptide and the fluorescent dye in our experimental extracellular vesicle analysis (*see experimental* *section 2.4*). Snapshots taken at the 50^th^ ns and 75^th^ ns during the 100 ns MD simulation were used as the input structures for the PepBD design algorithm.

The PepBD algorithm was next employed to design peptide sequences that bind to EpCAM using the GSG-EP-1-EpCAM NTD and EP-2-GSG-EpCAM P1 as the starting structures. For peptides to be effective as biosensors, they need to have specific properties such as good water solubility, appropriately-charged residues to strengthen electrostatic interactions with the target receptor, and structural stability. PepBD^24–27^ incorporates this knowledge by categorizing the 20 naturally-occurring amino acids into five types (hydrophobic, hydrophilic, positive charge, negative charge, and glycine). Each category represents a specific “hydration property”. The number of amino acids belonging to each type of hydration is fixed during the design process to improve peptide stability and increase the likelihood that the peptide will bind to the target. For example, the number of hydrophilic residues is fixed to enhance the likelihood that the peptide will be soluble, and the number of charged amino acids is fixed to complement the charges on the binding site of the protein. For our peptide design for EpCAM, four different hydration property cases were specified for the peptide chain. *N_H_, N_P_, N_-_, N_+_, N_O_, N_G_* each represent the number of hydrophobic, hydrophilic, negative, positive, other and glycine residues in the peptide sequence. **Case One:** *N_H_* = 4, *N_P_* = 3, *N_+_* = 0, *N_-_*= 2, *N_O_* = 0 and *N_G_* = 0, **Case Two:** *N_H_* = 4, *N_P_* = 3, *N_+_* = 1, *N_-_*= 1, *N_O_* = 0 and *N_G_* = 0, **Case Three:** *N_H_* = 4, *N_P_* = 2, *N_+_* = 1, *N_-_*= 2, *N_O_* = 0 and *N_G_* = 0, **Case Four** is: *N_H_* = 4, *N_P_* = 2, *N_+_* = 2, *N_-_*= 1, *N_O_* = 0 and *N_G_* = 0. Each *in-silico* search was performed for 10,000 Monte Carlo steps. As the search proceeded, new peptide sequences and conformers were generated, and the PepBD score (𝛤_score_) - a measure of the binding energy and the conformational stability of the peptide bound to the target protein - was recorded at each step. A lower 𝛤_score_ means stronger binding affinity of the peptide to the bound target (refer **Supplementary Figure S2** for PepBD score profile).

Once the *in-silico* PepBD evolution was completed, we performed explicit-solvent atomistic molecular dynamics (MD) simulations of the lowest-scoring peptides bound to EpCAM at the NTD and at the P1 pocket. We performed three 100 ns independent simulations for each of the lowest scoring peptide:EpCAM complex with the same initial coordinates but with different random velocities drawn from a Gaussian distribution. The Δ𝐺_𝑏𝑖𝑛𝑑𝑖𝑛𝑔_of the peptide:receptor complex at the end of the MD simulations was evaluated by using the MM/GBSA protocol and variable dielectric constant method. Table 2 presents the top two peptides, EP-2.1 and EP-2.2, identified from the EP-2-GSG-EpCAM P1 complex and the top two peptides, EP-1.1 and EP-1.2, derived from the GSG-EP-1-EpCAM NTD complex based on their binding free energy values. The table also includes their PepBD scores (𝛤_score_), binding free energy values (Δ𝐺_𝑏𝑖𝑛𝑑𝑖𝑛𝑔_) and solubility scores calculated from the CamSol Intrinsic method^47^ along with the peptide sequences. The CamSol Intrinsic method assesses the solubility profile for each residue in the peptide sequence and computes an overall solubility score. CamSol scores greater than 1 indicates highly-soluble peptides, while CamSol scores less than -1 denote peptides that prefer aggregation. The binding complexes formed by the designed peptides—EP-2.1 (sequence: ISIRNWEQW), EP-2.2 (sequence: EHRIRWWLN), EP-1.1 (sequence: WFMEENRQW), and EP-1.2 (sequence: WDDYTNFRY)—are depicted in **Figure 2**. Additional peptides for EpCAM identified during the design process are listed in Supplementary Table S1. Peptides EP-2.1, EP-2.2, EP-1.1 and EP-1.2 were selected for further experimental validation.

**Figure 2.**
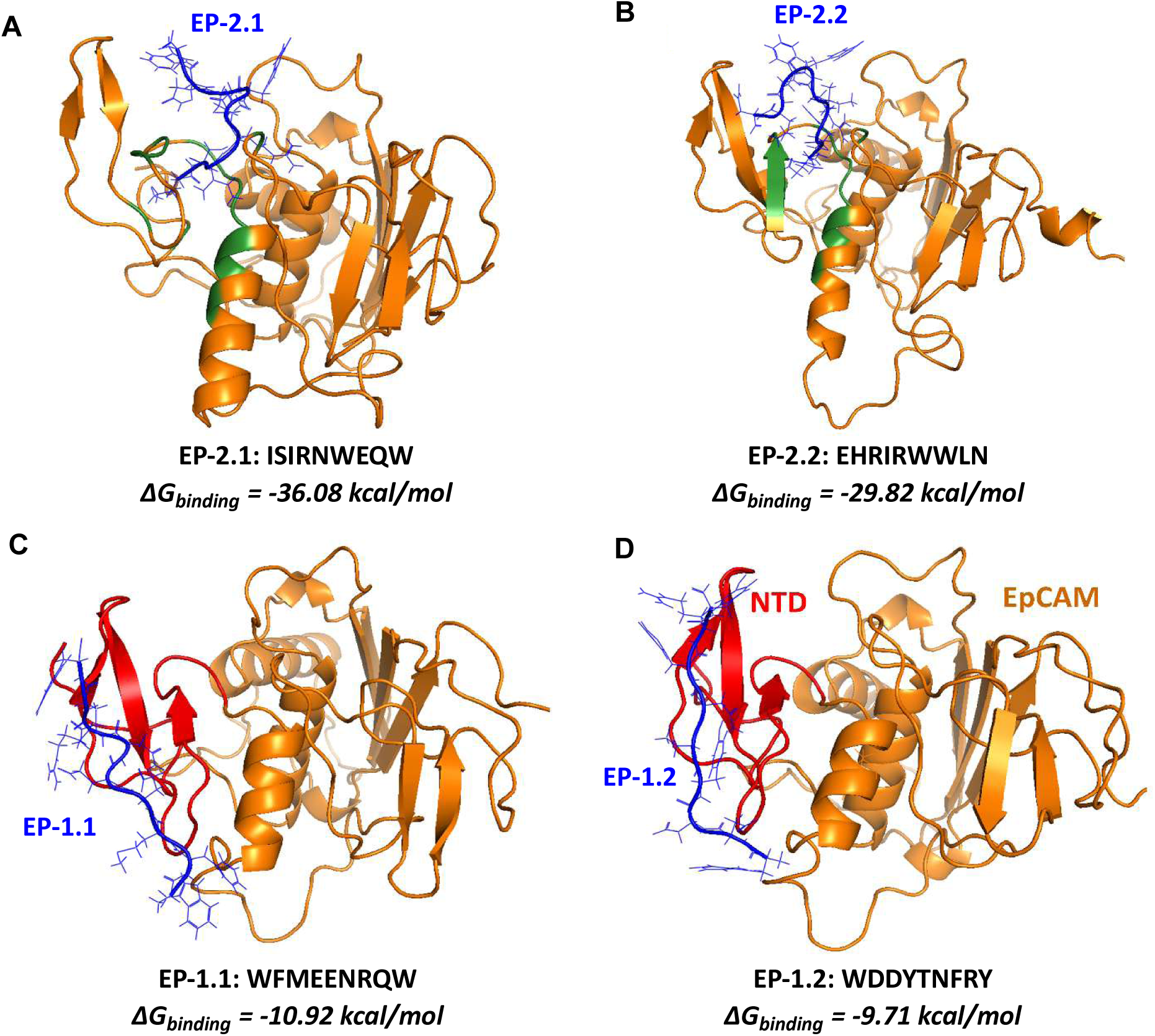
Snapshots of peptides **(A)** EP-2.1 (sequence: ISIRNWEQW), **(B)** EP-2.2 (sequence: EHRIRWWLN), **(C)** EP-1.1 (WFMEENRQW) and **(D)** EP-1.2 (sequence: WDDYTNFRY) bound to EpCAM obtained from molecular dynamics simulations.

**Table 2.** List of best peptide sequences identified by PepBD screening from EP-2 and EP-1 as the reference peptides with their corresponding CamSol solubility scores, 𝛤_score_ and Δ𝐺_binding_values. These peptide sequences were selected for synthesis and experimental validation.

| *Hydration*<br>*Property Case* | *Peptide* | *Peptide*<br>*Sequence* | *Solubility*<br>*Score* | *$\Gamma_{score}$*<br>*(kcal /mol)* | *$\Delta G_{binding}$*<br>*(kcal/mol)* |
| --- | --- | --- | --- | --- | --- |
| Reference<br>peptides | EP-2 | YEVHTYYLD | 0.77 | - | -6.11 |
|  | EP-1 | VSVHTYDLE | 0.74 | - | -2.41 |
| Case 2 | EP-2.1 | ISIRNWEQW | 1.72 | -56.55 | -36.08 |
| Case 4 | EP-2.2 | EHRIRWWLN | 1.32 | -64.35 | -29.82 |
| Case 3 | EP-1.1 | WFMEENRQW | 1.78 | -51.71 | -10.92 |
| Case 3 | EP-1.2 | WDDYTNFRY | 1.73 | -37.23 | -9.71 |

It is important to note that a low binding free energy (Δ𝐺_𝑏𝑖𝑛𝑑𝑖𝑛𝑔_) doesn’t always correspond to a low dissociation constant ***(K_D_)***, which needs to be determined experimentally. One reason for such discrepancies is that the MM/GBSA method used for calculating binding free energies, relies on an implicit-solvent model. This method may not fully account for the influence of the solvent, water, thus failing to capture the true enthalpy and entropy changes during desolvation for the target protein and the peptides upon binding. Furthermore, the MM/GBSA method assumes that the conformational states of the peptide ligand and protein receptor remain unchanged upon binding to form a complex. This assumption may not accurately reflect the dynamic nature of protein-ligand interactions, potentially leading to a discrepancy in the predicted binding affinity and the experimentally calculated dissociation constant^48,49^.

### 2.2 Computational design of peptides for CD81

#### 2.2.1 Generating initial structures for PepBD through docking simulations

As we did for EpCAM, we first determined the structure of the complex between reference peptide, P152 (CFMKRLRK), which was identified by Suwatthanarak et al.^36^, and our second target protein, the large extracellular loop (LEL or EC2) of tetraspanin CD81. The LEL domain of the tetraspanin proteins is known to be the binding region targeted by monoclonal antibodies and contains most of the protein-protein interaction sites^50^. Hence, we hypothesized that the entire LEL domain (PDB ID: 1G8Q^51^ ∼ 90 amino acids) could serve as a potential binding site for the peptides. The protein structure was refined and coordinate files for P152 were generated following the same protocol as was used for EpCAM.

We conducted docking simulations to identify favorable binding conformations of the reference peptide P152 to the LEL domain of CD81 using both Autodock Vina^52,53^ and HADDOCKv2.4^44,45^. Docking results were evaluated, and the top-scoring conformation from each software platform was further subjected to explicit-solvent molecular dynamics (MD) simulations. Free energies were also computed using the MM/GBSA method. The binding conformation from Autodock Vina had a Δ𝐺_𝑏𝑖𝑛𝑑𝑖𝑛𝑔_= -11.66 kcal/mol (**Figure 3A** and **3B**) and the binding conformation from HADDOCKv2.4 had a Δ𝐺_𝑏𝑖𝑛𝑑𝑖𝑛𝑔_= -9.29 kcal/mol (**Figure 3C** and **3D**). The two conformations showed similar interactions, with the two positively-charged residues on P152 interacting with the three negatively-charged residues on the LEL domain of CD81 (**Figure 3B** and **3D**). The binding conformation from Autodock Vina was selected for the peptide design because Δ𝐺_𝑏𝑖𝑛𝑑𝑖𝑛𝑔_was the lower of the two.

**Figure 3.**
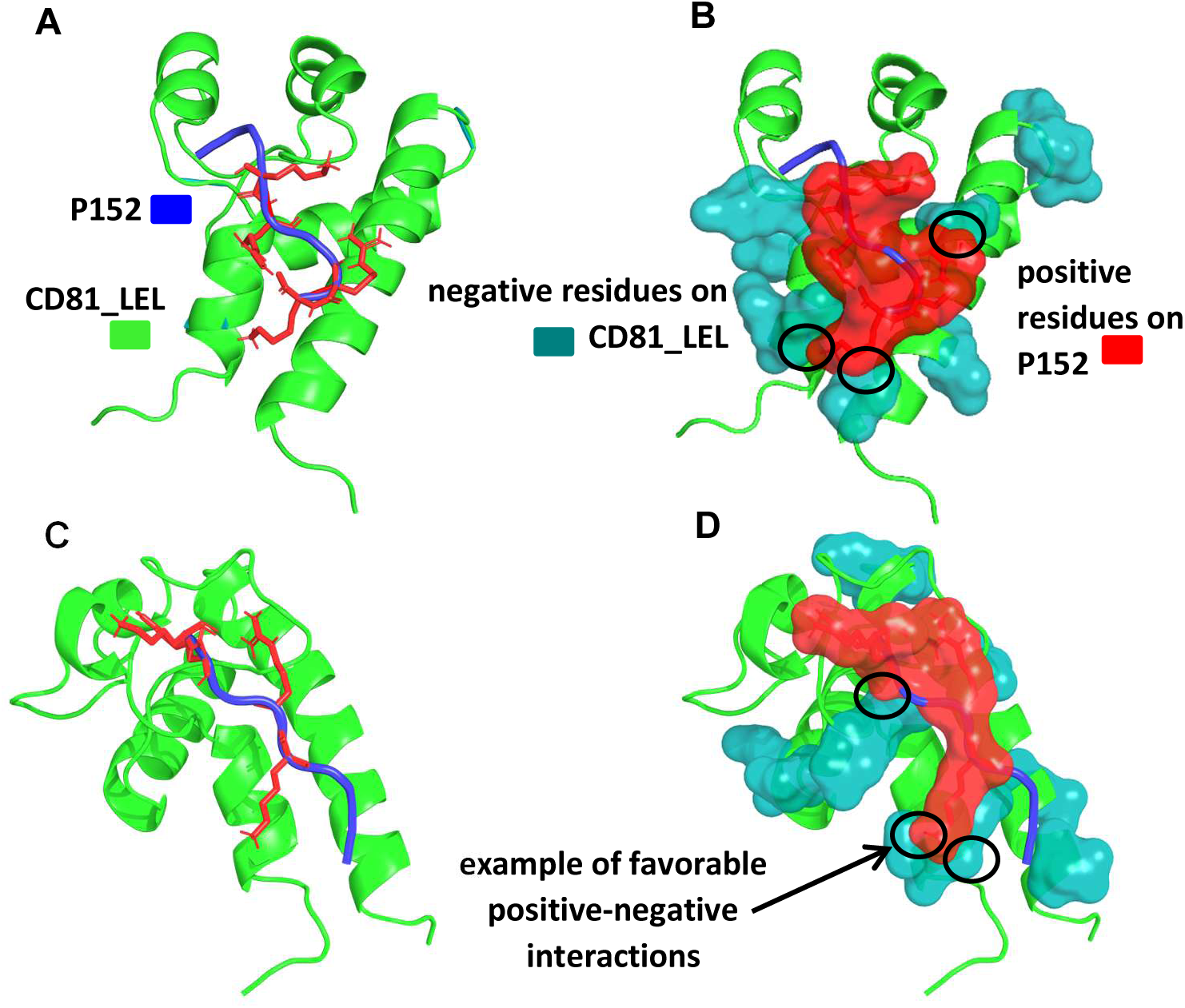
Best binding conformations between P152 and LEL domain of CD81 (CD81_LEL) (A) Best conformation of P152 bound to the CD81_LEL found through Autodock Vina with Δ𝐺_binding_ = -11.66 kcal/mol. This conformation serves as input for the first round of peptide design. (B) The same complex as in (A), highlighting favorable electrostatic interactions between the positively-charged residues 4 and 7 of P152 (shown in red) and the negatively-charged residues 5, 16, and 83 of CD81_LEL (shown in teal); these key interactions are circled in black. (C): Best conformation of P152 bound to the CD81_LEL found through HADDOCKv2.4 with Δ𝐺_binding_=-9.29 kcal/mol. (D) The same complex as in (C), showing key interactions (marked in black circles) between positively-charged residues 5 and 7 of P152 (red) and negatively-charged residues 77, 83, and 84 of CD81_LEL (teal).

#### 2.2.2 Design of Peptides to bind to the large extracellular loop (LEL) on CD81

##### First round of peptide design

Following the generation of the initial input structure from the docking simulations, we employed the PepBD algorithm to optimize the peptide sequence. The docking simulations revealed that the positively-charged residues on reference peptide P152 play a critical role in its binding interactions with the LEL domain of CD81 (CD81_LEL). Given this observation, we explored several design cases with varying hydration properties, primarily focusing on altering the net charge and the hydrophobicity of the sequences to design peptides suitable for biosensor applications. While reference peptide P152 lacked specific hydrophilic residues, we ensured that at least one hydrophilic residue was included in each design case to enhance the peptide solubility.

For the first design round, we performed 21 independent PepBD runs, using seven different sets of hydration property constraints, executing each three times for 10,000 MC steps. Each PepBD run started with a random trial peptide sequence and followed a unique search path. As the search commenced with random sequences, the initial PepBD scores tended to be high. However, as the peptide space was efficiently explored, multiple minima appeared in the score profile (refer to **Supplementary Figure S2**), indicating peptides with potentially-favorable binding affinities for the target protein. Supplementary Table S2 lists the best-scoring 8-mer peptides identified in each hydration property case. The top-scoring peptides from the initial design round were predominantly positively charged. Since the cell membrane possesses an overall negative charge^54,55^, there was a concern that these positively charged peptides could lead to non-specific binding with the cell membrane, instead of with the intended target protein site which was the LEL domain of CD81 (CD81_LEL). To address this, a second design round of peptide design was initiated aiming for peptides that were negative or neutral in charge.

##### Second round of peptide design

The top-performing sequence based on its low PepBD score with a net charge of +1 from the first round of design was peptide DQWLRARW (see Supplementary Table S2). However, because a positively-charged peptide could lead to non-specific interactions with the negatively-charged cell membrane, we initiated a second round of design to identify peptides with a neutral or negative charge to minimize such effects. We modified DQWLRARW by introducing one glutamic acid (E) and one aspartic acid (D) at different positions, creating four new reference peptides CD81-1, CD81-2, CD81-3 and CD81-4 (Table 3), each with a net charge of -1. Since the original peptide’s binding relied on positively-charged residues interacting with negatively-charged residues on CD81_LEL, adding negative residues disrupted these interactions, necessitating a new search for optimal binding sites. Following this modification, docking simulations for the new reference peptides were performed using AutoDock Vina and HADDOCKv2.4, as in the previous round. To further refine the best-scoring docking conformations of each new reference peptide with CD81_LEL, 100 ns molecular dynamics (MD) simulations were conducted, followed by binding free energy calculations.

**Table 3.** New reference peptides (CD81-1, CD81-2, CD81-3 and CD81-4) generated by adding negative residues (in green) at different locations to the peptide DQWLRARW, which was obtained from the first round of design. Results from free energy calculations performed using the MM/GBSA method for the binding conformations obtained from HADDOCKv2.4 and Autodock Vina for each new reference peptide.

| *New reference peptides generated from DQWLRARW* | *Energy values calculated using MM/GBSA for binding conformation found through docking (kcal/mol)* |  |
| --- | --- | --- |
|  | *Autodock Vina* | *HADDOCKv2.4* |
| <u>DD</u>QWLRARW<u>E</u><br>(CD81-1) | 3.99 | 0.81 |
| DQWL<u>DE</u>RARW<br>(CD81-2) | <u>-0.67</u> | <u>-8.08</u> |
| DQWELR<u>D</u>ARW<br>(CD81-3) | 2.09 | <u>-17.91</u> |
| DQ<u>E</u>WLRA<u>D</u>RW<br>(CD81-4) | 3.73 | 0.66 |

Binding conformations of the new reference peptides (CD81-1, CD81-2, CD81-3 and CD81-4) with the CD81_LEL domain that exhibited negative free energies were identified as plausible starting input structures for the second set of PepBD runs. These favorable free energies are highlighted as bold and underlined in Table 3). Among these, Autodock Vina produced only one favorable case of negative binding free energy for the conformation between the new reference peptide CD81-2 and CD81_LEL with Δ𝐺_𝑏𝑖𝑛𝑑𝑖𝑛𝑔_= -0.67 kcal/mol. In contrast, HADDOCKv2.4 identified two favorable interactions: CD81-2 binding to CD81_LEL with a Δ𝐺_𝑏𝑖𝑛𝑑𝑖𝑛𝑔_= -8.08 kcal/mol and CD81-3 binding with CD81_LEL with a Δ𝐺_𝑏𝑖𝑛𝑑𝑖𝑛𝑔_= -17.91 kcal/mol (Table 4). The binding conformations from HADDOCKv2.4 were selected for the second round of peptide design due to their more favorable (lower) free energy values. Additionally, the binding locations from HADDOCKv2.4 were further away from the cell membrane (**Figure 4A**) than the binding sites predicted from Autodock Vina (**Figure 4B**), thus reducing the likelihood of non-specific interactions with the cell membrane.

**Figure 4.**
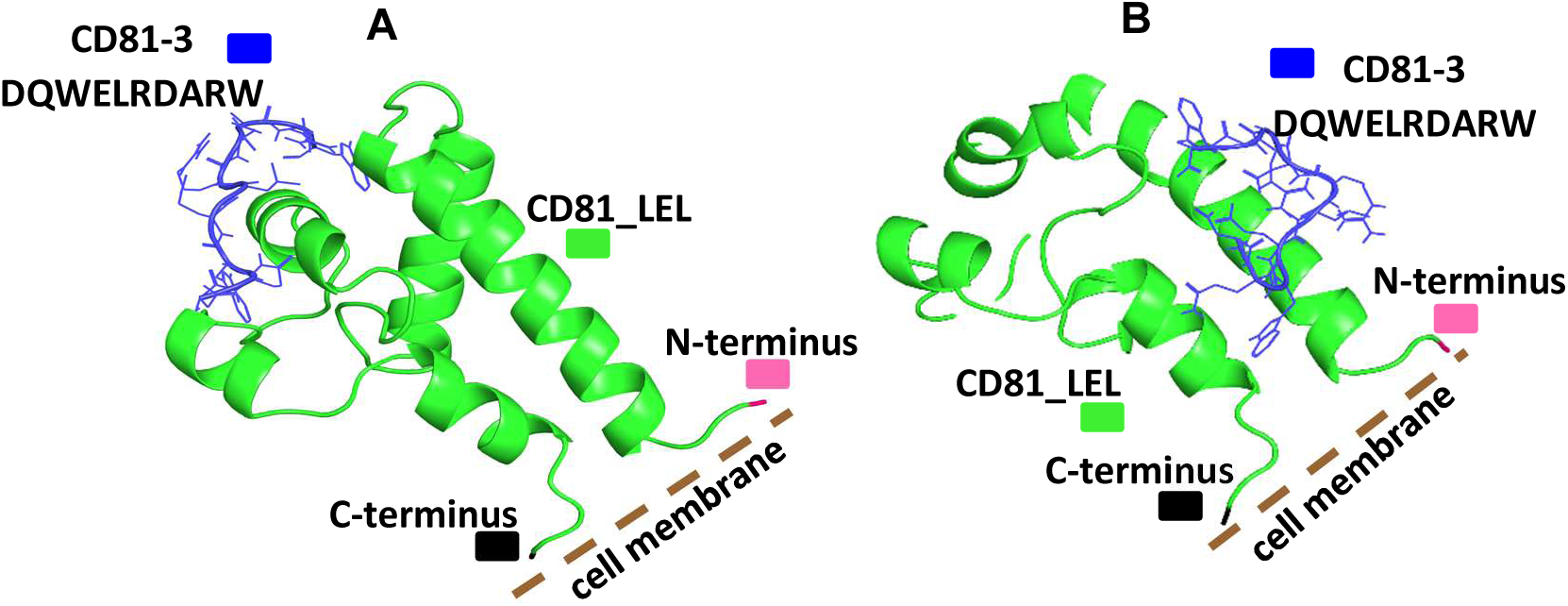
Binding conform erence peptide CD81-3. (A) obtained from HADDOCKv2.4 and **(B)** from Autodock Vina. The binding site identified from Autdock Vina is closer to the cell membrane (represented as a dotted brown line) than the binding site identified from HADDOCKv2.4.

**Table 4.**
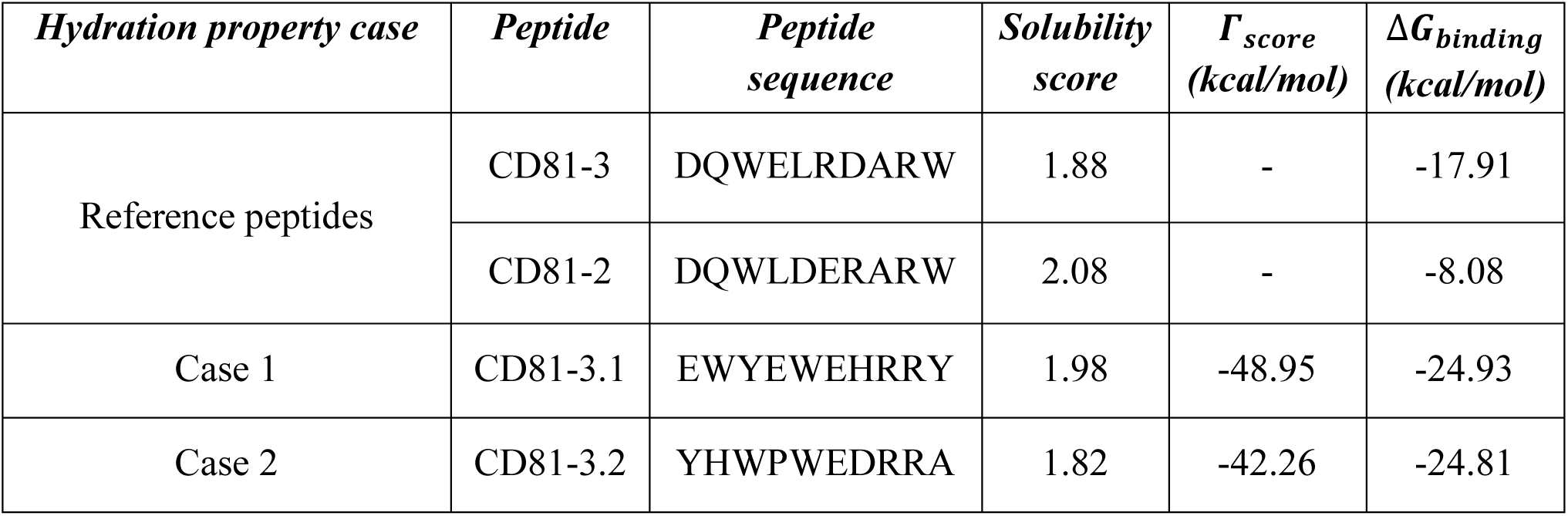
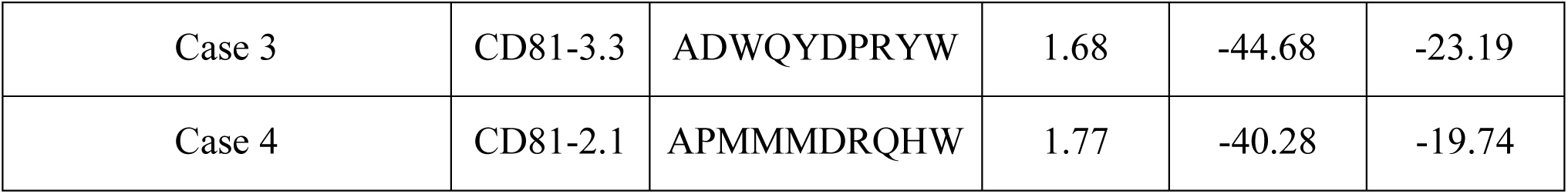
Best computationally designed peptide ligands for the LEL domain of CD81. The peptide sequences are listed along with their PepBD scores, intrinsic solubility scores as calculated through the CamSol method, and best binding free energies (Δ𝐺_𝑏𝑖𝑛𝑑𝑖𝑛𝑔_in kcal/mol) computed from running three independent explicit-solvent MD simulations for 100ns. The specifications for the hydration property cases are provided in Supplementary Table S3. These peptide sequences were selected for synthesis and further experimental validation.

For the second round of peptide design, we focused on exploring different hydration property combinations to design peptides with negative or neutral net charge. Binding conformations from HADDOCKv2.4 for reference peptides CD81-2 and CD81-3, were used as the starting points for further optimization. Each reference peptide underwent PepBD runs for seven hydration property combinations, with three runs per combination using different random seeds. Therefore, a total of 42 independent PepBD run (7 design cases × 3 random numbers × 2 reference peptides) were performed for the second round of peptide design. The lowest-scoring peptides from each PepBD run were then subjected to an explicit-solvent molecular dynamics simulation for 100 ns to assess their binding dynamics with the LEL domain of tetraspanin CD81. New hydration property design cases and the top 10 peptide candidates from the PepBD runs for CD81 are reported in Supplementary Table S3. A more negative value of free energy (Δ𝐺_𝑏𝑖𝑛𝑑𝑖𝑛𝑔_) indicates a stronger binding affinity between the peptide sequence and the target protein. Four peptide candidates, (presented in Table 4) CD81-3.1, CD81-3.2, CD81-3.3 and CD81-2.1, exhibited lower binding free energies than the reference peptides CD81-3 and CD81-2 making them suitable for further experimental validation. Solubility scores were also computed to predict the solubility of the discovered peptides using the CamSol Intrinsic method. Table 4 includes these solubility scores along with PepBD scores and their free energy values.

### 2.3. Peptide synthesis

After identifying effective anti-EpCAM and anti-CD81 peptide sequences, we chose eight peptides—EP-2.1, EP-2.2, EP-1.1, EP-1.2 for EpCAM, and CD81-3.1, CD81-3.2, CD81-3.3 and CD81-2.1 for CD81—for experimental testing due to their significant Δ𝐺_𝑏𝑖𝑛𝑑𝑖𝑛𝑔_ values. These peptides were synthesized at the MGH Peptide Core utilizing the Fmoc solid-phase method with an Apex 396 automatic synthesizer (AAPPTec, LLC).

We next performed the conjugation of fluorescent dyes to these peptides. Initially, we attached Fmoc-N-PEG-COOH to the NH2 group at the N-terminal of the peptides. In this procedure, we coupled the amino group (NH2) at the end of the peptide with the carboxylic group (COOH), using HOBt (1-Hydroxybenzotriazole), HBTU (O-Benzotriazole-N,N,N’,N’- tetramethyluronium hexafluorophosphate), and DIEA (N,N-Diisopropylethylamine) as reagents^56^. HOBt and HBTU activate carboxylic acids for enhanced reactivity with amines, while DIEA regulates pH and boosts activation when used with HBTU, effectively promoting amide bond formation. The carboxyl group was subsequently linked at the peptide’s end through an amide bond formation. Deprotection of the Fmoc group was achieved using piperidine, followed by the creation of an NHS ester-NH2 linkage (**Figure 5A**). The resultant 5[6]-FAM-PEG-modified peptides (termed F-peptides hereafter) were purified using HPLC^57,58^

**Figure 5.**
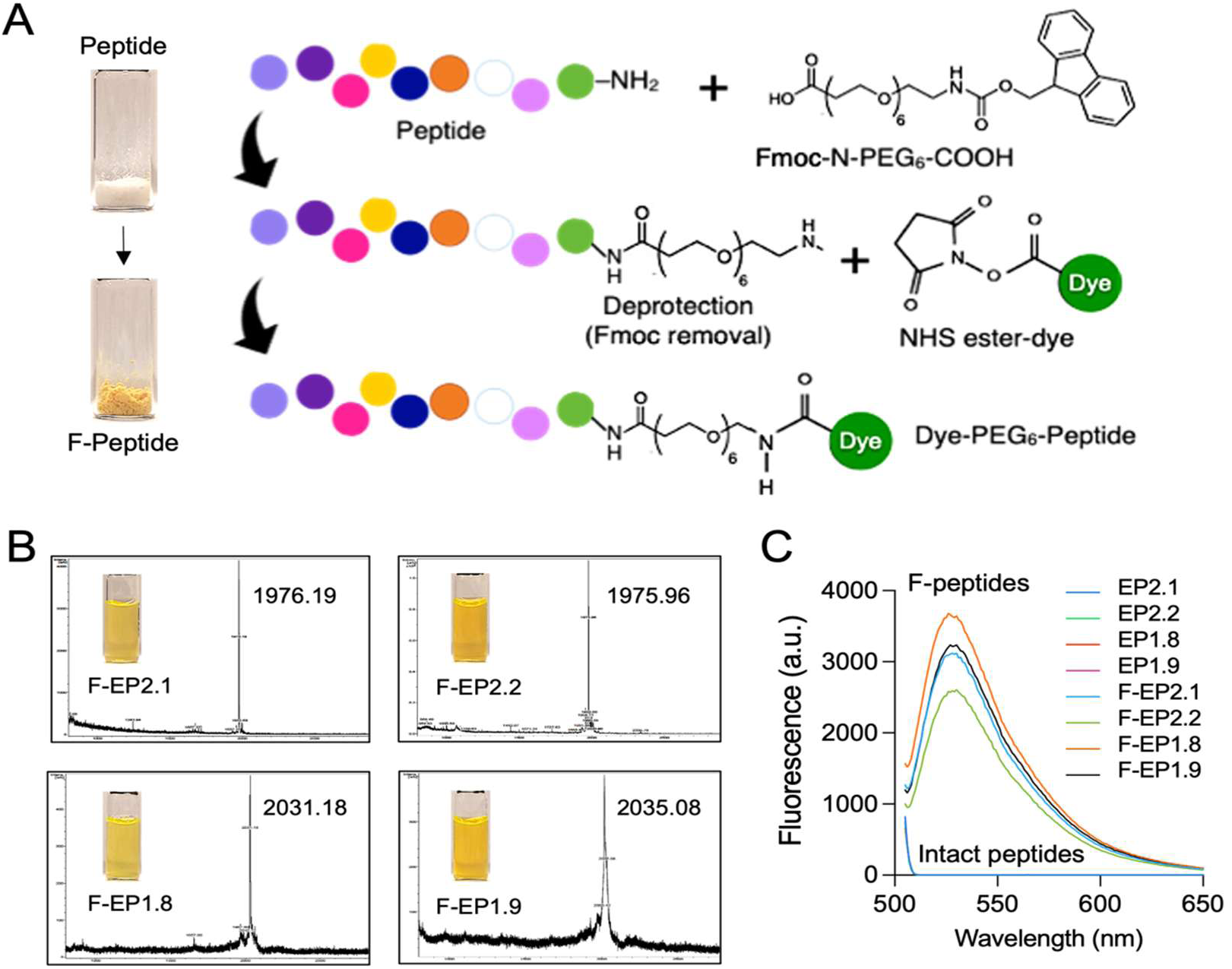
Preparation and characterization of 5[6]-FAM-PEG-modified peptides (F-peptide) (A) Schematic images of peptide modification with fluorescent dye to form an amide bond between amine and carboxylic acid groups utilizing HOBt, HBTU, and DIEA. For dye conjugation, we employed NHS ester-dye, which was attached to the N-PEG6 group. (B) Measurement of MALDI-MS to validate the F-peptides molecular weights. A significant single peak indicating the exact molecular weight of F-peptides appeared. (C) Fluorescence measurement. All F-peptides generated strong fluorescence around 525 nm with 488 nm excitation

Post-PEG modification, we coupled amine-reactive fluorescent dyes, NHS-Alexa-488, to PEG via NH2-NHS chemistry and purified the dyed peptides using HPLC. The synthesized peptides underwent validation through matrix-assisted laser desorption/ionization mass spectrometry (MALDI-MS, **Figure 5B**). The four F-peptide candidates targeting EpCAM showed peaks at their theoretically expected mass: F-EP-2.1 (1976.19), F-EP-2.2 (1975.96), F-EP-1.1 (2031.18), and F-EP-1.2 (2035.08). Similar results were observed for the F-peptide candidates for CD81: F-CD81-3.1 (2249.22), F-CD81-3.2 (2111.42), F-CD81-3.3 (2095.07) and F-CD81-2.1 (2042.56). We used UV-visible spectroscopy to illustrate the verification of the fluorescent dye presence in the peptides (**Figure 5C**). Under 488 nm light excitation, these peptides exhibited strong fluorescence at 530 nm, confirming the successful labeling with 5[6]-FAM, unlike the non-fluorescent intact peptides.

### 2.4. Validation of the 5[6]-FAM-PEG-modified peptide’s (F-peptides) affinity for the target proteins. EpCAM and CD81

We assessed the performance of F-peptides in detecting EpCAM and CD81, (**Figures 6** and **7**). Initially, we conducted a standard fluorescent ELISA. Briefly, various concentrations of EpCAM (ranging from 1 to 1,000 ng/mL) and CD81 (from 0.1 to 100 ng/mL) were independently adsorbed onto a 96-well plate. To evaluate the non-specific binding of peptides, non-targeting control peptides (CP1:QWWIRNEIS, CP2:NEHRLRWIW, CP3:QFRMWEENW, and CP4:DYTWDRNYF), which contain the same amino acids as the designed peptides (Table 2 and 4) but arranged in a scrambled order, were used. For EpCAM, peptides F-EP-2.1 and F-EP-1.2 exhibited increasing fluorescence signals with rising EpCAM concentrations (**Figures 6A** and **6D**), indicating strong affinity for the target protein. In contrast, F-EP-2.2 and F-EP-1.1 showed weak binding. Similarly, for CD81, peptides F-CD81-3.1 and F-CD81-3.3 demonstrated enhanced fluorescence signals as CD81 concentrations increased (**Figures 7A** and **7C**), suggesting favorable affinity for the protein. The control peptides, with their scrambled sequences, yielded only minimal fluorescent signals. We chose F-EP-2.2 (for EpCAM) and F-CD81-3.1 (for CD81) for further single EV imaging validation, based on their generation of high signals in response to respective target proteins.

**Figure 6.**
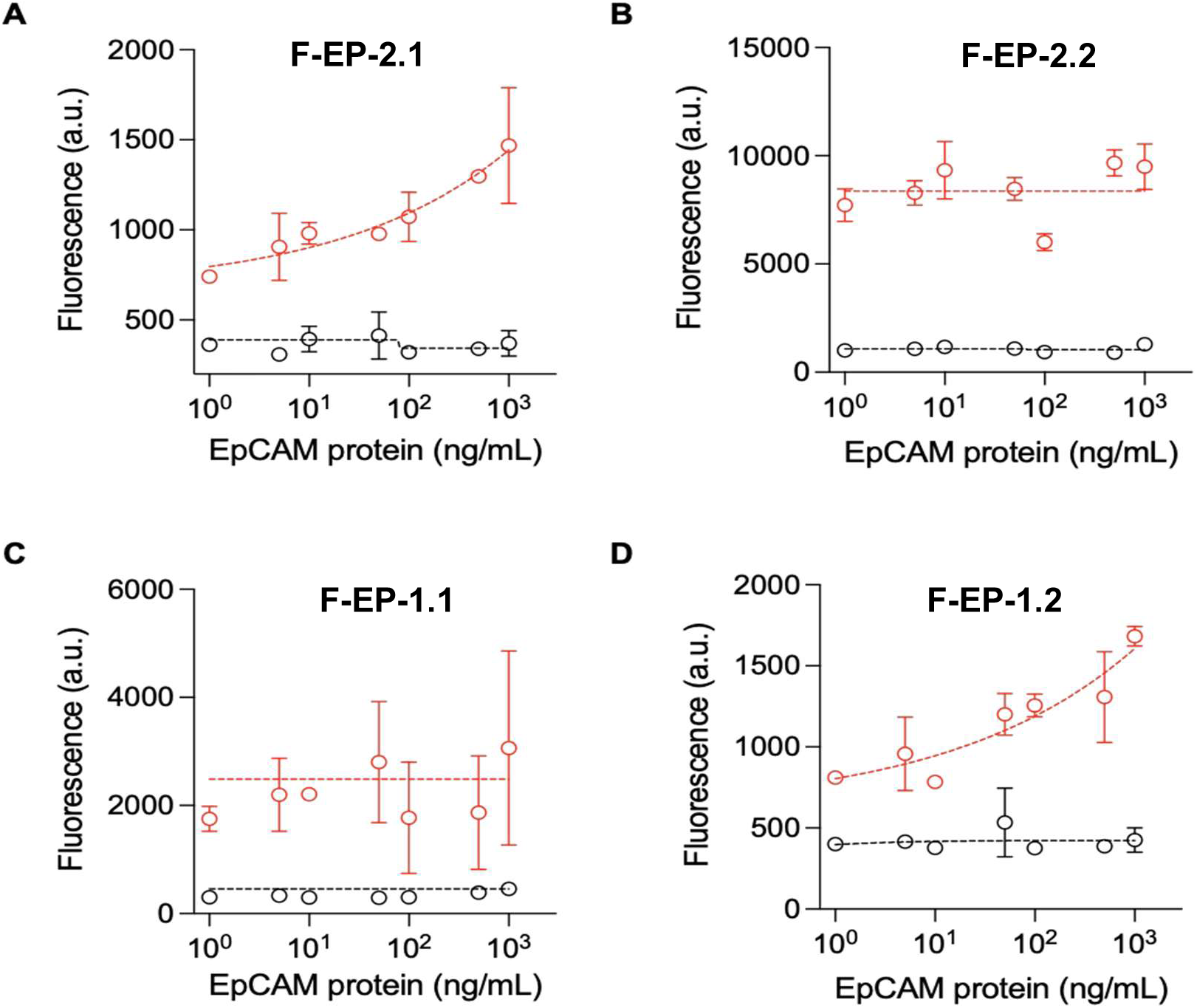
Validation of the EpCAM F-peptides. The affinity of the EpCAM F-peptide for EpCAM protein was assessed using a fluorescent ELISA. We labeled fluorescent dye (488 nm) on the peptide. In the figure, the red color indicates the signal from the EpCAM peptide, while the black color represents the control peptide signal. (A) EP-2.1(B) EP-2.2 (C) EP-1.1 and (D) EP-1.2 peptides. The data are presented as means ± standard deviation from measurements taken in duplicates.

**Figure 7.**
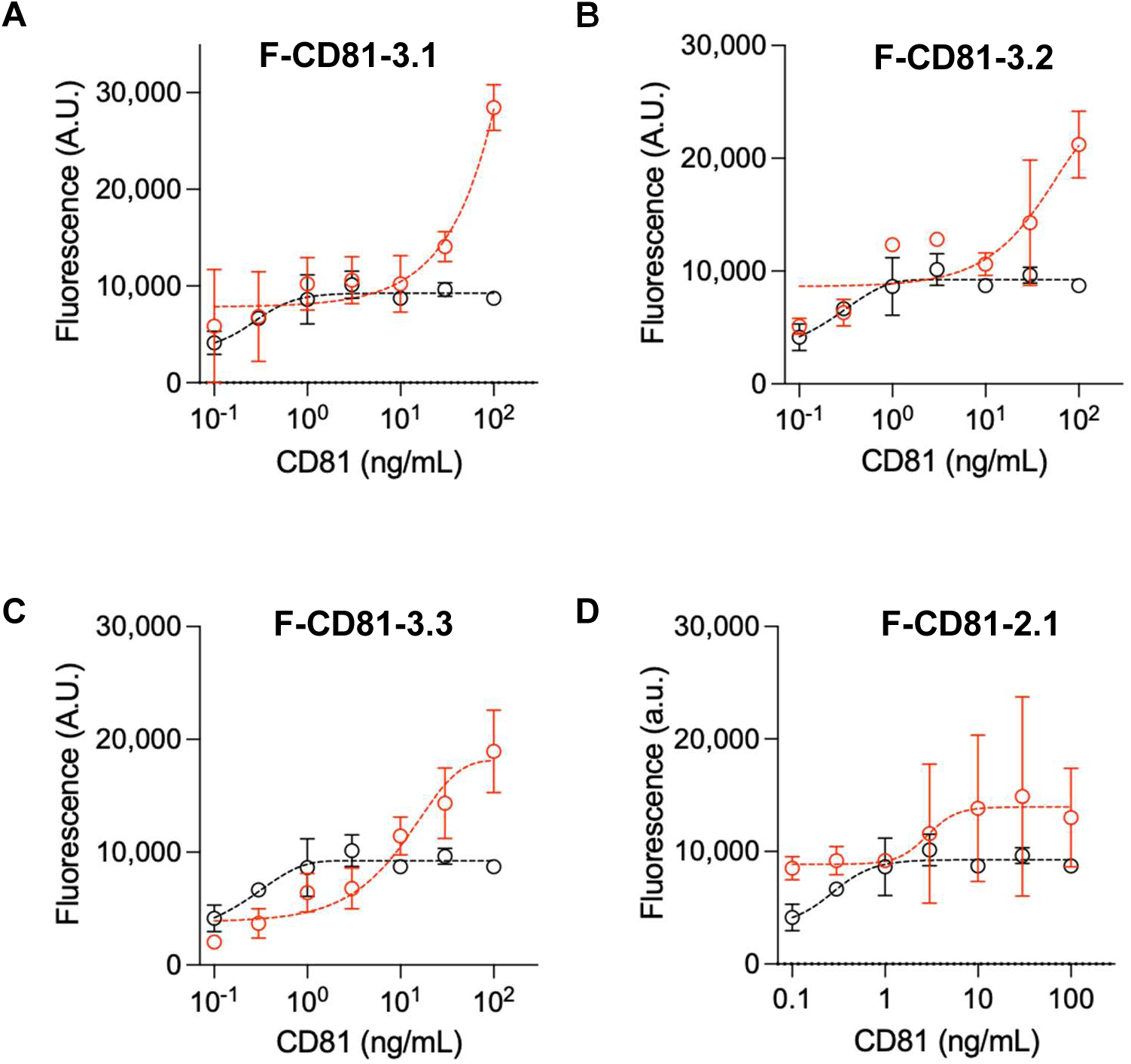
Validation of the CD81 F-peptides. We used a fluorescent ELISA to evaluate CD81 peptide affinity to the CD81 protein. The CD81 F-peptide signals are represented in red, while the control peptide signals are in black. (A) CD81-3.1, (B) CD81-3.2, (c) CD81-3.3 and (D) CD81-2.1. The results are shown as means ± standard deviation from measurements conducted in duplicates.

### 2.5 Single EV imaging analysis

We then proceeded with single EV imaging **(Figure 8)**. EVs were extracted from SW620, MCF7, and MDA-MB-231 cells using a size exclusion chromatography (SEC) column. The cell medium was first concentrated using a Centricon Plus-70 Centrifugal filter and then centrifuged. This concentrated medium was subsequently processed through SEC, and the 4th and 5th fractions were collected for EV isolation. Initially, the EV membranes were stained with Alexa-555 dye to pinpoint the localized EVs on a substrate. Stained EVs were subsequently incubated with the F-peptides, either targeting EpCAM or CD81 on the EV surface **(Figure 8A)**. We used peptide candidates validated in ELISA as imaging probes: F-EP-2.1 for EpCAM **(Figure 6)** and F-CD81-3.1 for CD81 **(Figure 7)**. Labeled EVs were imaged via confocal microscopy. The fluorescent signal at 565 nm (red dot) was used to locate EVs, and the signal at 515 nm (green dot) was used to measure the protein target (EpCAM or CD81). To compare with F-peptide imaging, we prepared a separate set of EV samples using fluorescent antibodies (F-antibody) for EpCAM or CD81 labeling.

**Figure 8.**
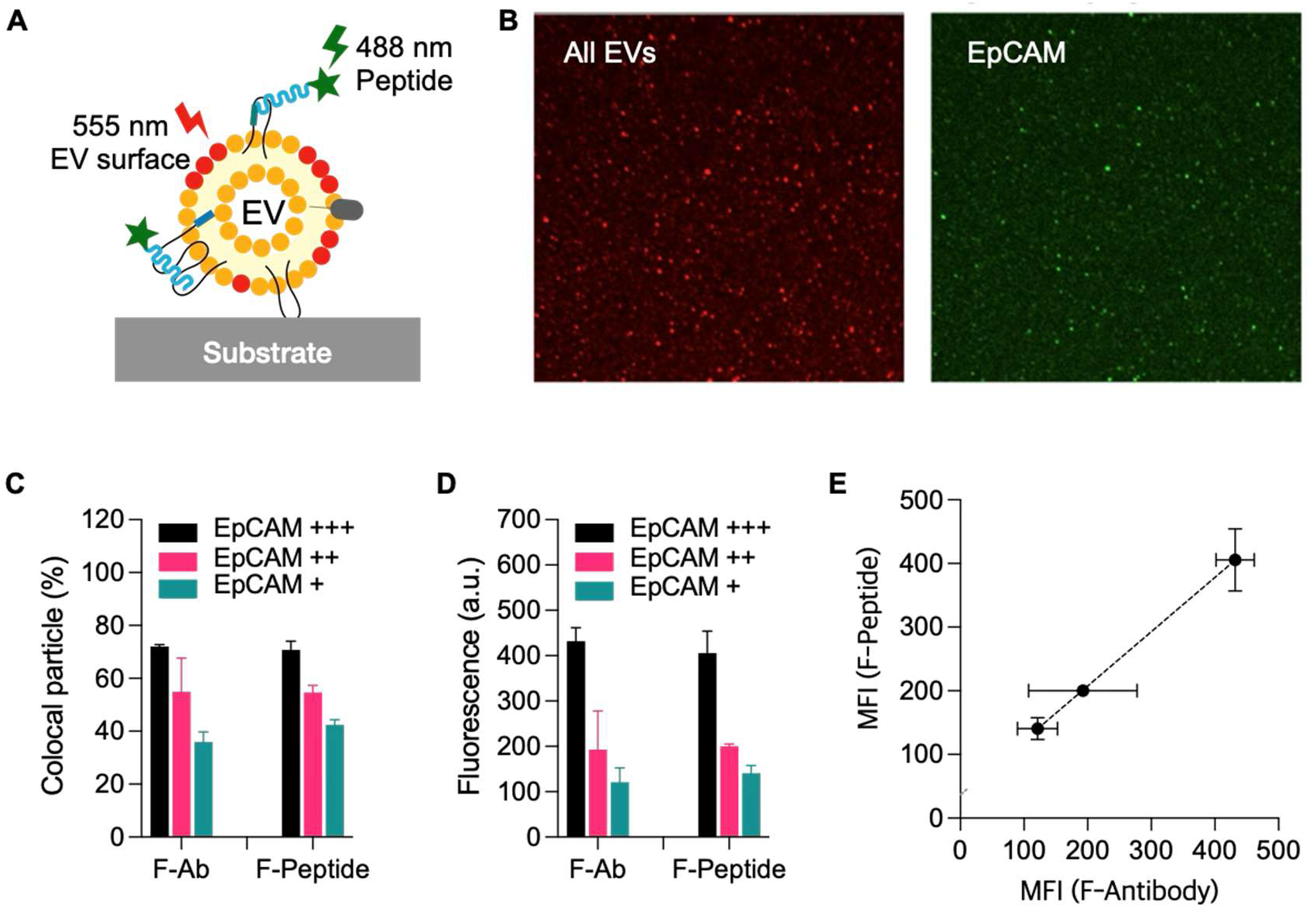
Single EV analysis for EpCAM peptide (A) Schematic image of single EV imaging. Different fluorescent dyes, Alexa-555 (red) and 5(6)-FAM (green) were labeled at the EV membrane and the developed peptide respectively. Fluorescence images were first taken from all EVs (in red) and after F-peptides were added (green). (B) Microscope images. Red dots represent EVs and green signal came from EpCAM peptide F-EP-2.1,which was able to detect EVs. (C) Spatial overlap (colocalization) of red (EV) and green (EpCAM) signals. The results confirmed that F-EP-2.1 (green) bound to EVs (red). This suggests that the green signals are not due to nonspecific binding to the substrate (glass), but rather to EpCAM on the surface of the EVs. Peptide and antibody showed similar affinity to target EpCAM on the EVs. (D) Fluorescence of F-antibody and F-peptide, depending on different EpCAM expressions (high:+++, moderate:++, low:+), are plotted. Both values corresponded well to the expression rate. (E) MFIs (mean fluorescent intensities) of antibody- and peptide-based EV imaging were comparable (R^2^=0.967).

**#Figure 8B** presents the EV imaging results for EpCAM detection. Both F-peptide and F- antibody exhibited a similar performance. We observed differential signals in EpCAM-positive EV counts **(Figure 8C)** and fluorescent intensities **(Figure 8D)** across cell lines exhibiting varying EpCAM expression levels: low (+, MDA-MB-231), moderate (++, MCF7), and high (+++, SW620). Conversely, EV counts and fluorescent intensities were comparable between peptide and antibody probes when analyzing EVs derived from identical cell lines. Consequently, the mean fluorescent intensities (MFIs) showed a strong linear correlation (*R*^2^ = 0.967) between peptide- and antibody-based EV imaging **(Figure 8E)**. A comparable trend was observed in the CD81 imaging results **(Figure 9)**, where a strong correlation (*R*^2^ = 0.998) was demonstrated between the imaging data obtained using the F-peptide and F-antibody probes.

**Figure 9.**
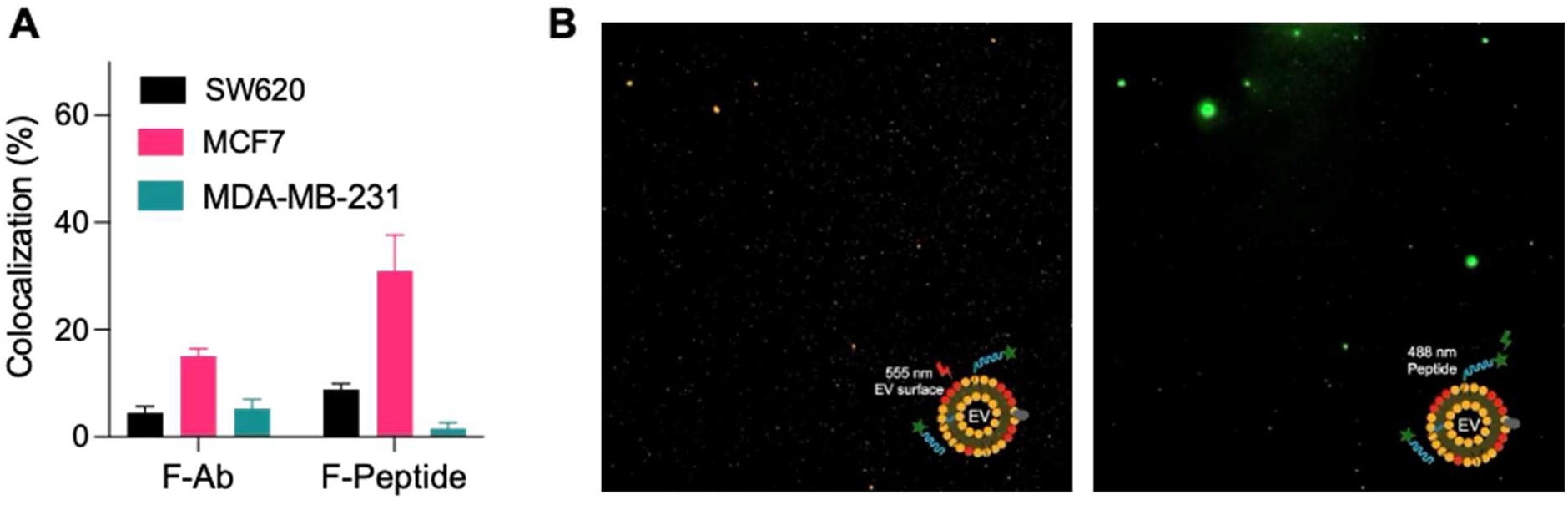
Single EV analysis for peptide F-CD81-3.1 with EVs extracted from three different cell lines SW620, MCF7 and MDA-MB-321. (A) Colocalization results indicated that both the peptide and the antibody demonstrated comparable affinities for the target CD81 on the EVs. (B) Microscope images showed red dots (555 nm) representing EVs, with green signals (488 nm) originating from the CD81 peptide CD81-3.1.

## 3. Conclusion and Future Work

Peptide-based biosensors offer a promising approach for detecting extracellular vesicles (EVs) through biomarkers such as tetraspanins. Moreover, they can be adapted to identify EVs originating from cancerous cells, providing potential applications in disease diagnostics. This study integrates computational and experimental methodologies to design peptide ligands capable of capturing extracellular vesicles, specifically exosomes—by targeting tetraspanin CD81, a transmembrane protein on exosomes. Additionally, we sought to distinguish cancerous from non-cancerous exosomes by developing peptide ligands that selectively bind to EpCAM, a protein overexpressed in cancer cells. By employing the PepBD algorithm alongside computational techniques such as docking simulations, explicit-solvent molecular dynamics, and MM/GBSA calculations, we identified multiple novel peptide sequences predicted to have lower binding free energies than the reference peptides EP-1 (YEVHTYYLD) and EP-2 (VSVHTYDLE) for EpCAM (previously identified by Weizhi et al.) and P152 (CFMKRLRK) for CD81 (previously identified by Suwatthanarak et al.). Based on these findings, we selected eight promising peptide sequences—EP-2.1, EP-2.2, EP-1.1, and EP-1.2 for EpCAM, as well as CD81-3.1, CD81-3.2, CD81-3.3 and CD81-2.1 for CD81. These peptides were synthesized and first validated through standard fluorescent ELISA. Based on their signal strengths, peptides EP-2.2 (for EpCAM) and CD81-3.1 (for CD81) were selected for single EV imaging analysis. The peptide probe EP-2.1 performed comparably to the antibody probe in identifying EpCAM-positive EVs, as demonstrated by the mean fluorescent intensities (MFIs), which revealed a high correlation (R² = 0.967) between peptide- and antibody-based EV imaging. Similarly, the CD81-targeting peptide CD81-3.1 exhibited equivalent binding behavior, showing an even stronger correlation with the antibody probe (R² = 0.998) in EV imaging.

In future work, we aim to employ the methods used in this study to develop a peptide-based library that can target – 1) other EV-related markers such as TSG101 and Alix along with the designed probes for CD81, which will be instrumental in identifying and quantifying of extracellular vesicles and 2) tumor biomarker CD24 which when combined with EpCAM will improve the capture of cancerous EVs. Additionally, we plan to integrate these probes onto carrier substrates, such as nanoparticles. Due to their smaller size compared to antibodies, fluorescently labeled peptide probes can be more densely packed on these surfaces, increasing their availability and potency. This approach is expected to enhance immunoassay signal strength and improve targeting efficiency.

## 4. Methods and Materials

### 4.1 Computational Peptide Design

The **Pep**tide **B**inding **D**esign (PepBD) algorithm^24–27^ is a Monte Carlo-based method that facilitates the exploration of both the conformational and sequence space of peptides given a fixed number of amino acids in the peptide sequence. Through an iterative process, it identifies optimized peptide sequences that bind with higher affinity to a biomolecular target compared to a previously identified “reference peptide”. A detailed description and process flow of the PepBD algorithm can be found in our previous works.

The first step involves generating input complexes between a known peptide ligand and the target proteins. In this case, reference peptides EP-2 and EP-1 for EpCAM and P152 for CD81 were used. This step is independent of PepBD and utilizes docking platforms such as HADDOCKv2.4^44,45^ and AutoDock Vina^52,53^ to perform docking simulations. The reference peptides were modeled as flexible to allow backbone conformational changes, facilitating the exploration of optimal binding poses with the target protein. The docking results were then evaluated using molecular dynamics (MD) simulations and binding free energy values were calculated using the MM/GBSA protocol^39^. Binding conformations with the most favorable free energy values were selected as inputs for the peptide optimization process within PepBD.

The next step focuses on peptide optimization. Within the PepBD algorithm, random modifications, referred to as “trial moves,” are introduced in either the peptide sequence or its conformation. A random number (R) is generated and compared to a user-defined sequence change probability (P_sequence_). If R is less than or equal to P_sequence_, a sequence change occurs, while if R is greater than P_sequence_, a conformational change is initiated.

Sequence modifications follow one of two approaches - the first approach is an amino acid mutation, where a residue is replaced with another amino acid of the same hydration property type. The second approach is an amino acid exchange, where two amino acids swap positions within the sequence, regardless of their type. Similarly, conformational modifications can occur through three ways - first either the N-terminus or C-terminus rotates along with two residues in the middle of the sequence; second approach involves rotation of the entire peptide backbone, using both the N- and C-termini along with a single middle residue; third involves concerted rotation (CONROT) with three consecutive residues undergo displacement while keeping both termini fixed.

After each trial move, PepBD calculates a binding energy score (PepBD score) for the modified peptide. The new score is then compared to the previous score’s step, and the move is either accepted or rejected based on the Metropolis criterion. This constitutes one PepBD cycle, and the peptide design process runs iteratively for 10,000 such cycles.

At the end of the optimization process, the best peptide variants—those with the lowest PepBD energy scores—are selected for further analysis. These optimized peptide-protein complexes are then subjected to explicit-solvent atomistic molecular dynamics simulations to assess their binding affinity.

### 4.2 Atomistic Molecular Dynamics Simulation

Explicit-solvent atomistic MD simulations are an important tool to analyze the dynamics of the binding process between peptide and the target protein. All MD simulations are performed in the canonical (NVT) ensemble using the AMBER20 package^38^ to investigate the binding between the peptide sequences and the target proteins (EpCAM and CD81). The starting configurations of the peptide:protein complexes in each MD simulation were the output from the searches in the PepBD algorithm. This system was enclosed in a periodically truncated octahedral box of 12 Å and was explicitly-solvated in water by using the TIP3P water model^59^ within the AMBER20 package. Counterions such as Na+ or Cl- were added to neutralize the peptide:protein complex prior to running the MD simulations. Three independent simulations were carried out for each peptide:protein complex for 100 ns to ensure that the system reaches an equilibrated state. We used the implicit-solvent molecular mechanics/generalized Born surface area (MM/GBSA) approach with the variable internal dielectric constant model^46^ for post-analysis of the last 5 ns simulation trajectories of the peptide:protein complexes to evaluate the binding free energies. Details of post-analysis of the atomistic MD simulations can be found in our previous work.

### 4.3 Peptide synthesis

Fmoc-L-Ile -2-chlorotrityl resin comprising of the first amino acid of the peptide in C-terminal was preloaded on an automatic peptide synthesizer (Apex 396 model, AAPPTec, LLC). O- (benzotriazol-1-yl)-N,N,N0,N0 -tetramethyl uronium hexafluoro phosphate and thioanisole, 1- hydroxybenzotriazole were used as peptide coupling and activating agents. Diisopropylethylamine was used as the neutralizing reagent in three-fold molar excess for three-fold molar excess of Fmoc-amino acids to activate the carboxyl group of the incoming amino acid. Before the removal of the resin, Fmoc was removed from the N-terminus of the assembled peptide using 20% (v/v) of piperidine in dimethyl formamide (DMF). Subsequently, the resin was washed with DMF, dichloromethane (DCM), and isopropanol (IPA). A concoction of cleaving, as well as scavenging agents, was prepared with the following composition TFA/ethanedithiol/water/phenol/thioanisol in a ratio of 8:0.25:0.5:0.75:0.5 (v/v) to cleave the peptide from chlorotrityl resin and remove the side-chain protecting groups in 50% TFA in DCM (v/v)/ethanedithiol/water/phenol/thioanisol 8:0.25:0.5: 0.75:0.5 (v/v) for 1 h at RT. The resin was filtered, and ice-cold diethyl ether was added to extract the crude peptide precipitate and washed thrice with ice-cold diethyl ether. The residual ether was removed under pressure and the peptide was HPLC purified.

### 4.4 Peptide modification with fluorescent dye

The synthesized intact peptides were labeled with 5[6]-carboxy-fluorescein (5[6]-FAM). The protection group, the Fmoc, was removed by mixing with 20%(v/v) of piperidine in DMF. After sequential washing with DMF, DCM, and IPA, NHS-5[6]-FAM was reacted to the exposed N-terminus of the intact peptides in a dark room at RT. The fluorescent-modified peptides were then purified by HPLC.

### 4.5. Mass spectrometer measurement

All the peptides were analyzed by MALDI-MS on a Bruker Microflex instrument. Samples were co-crystallized with alfa-cyano-4-hydroxycinnamic acid and dried on the MALDI-MS plate at RT. Positive as well as negative modes of ionization were utilized.

### 4.6. Fluorescence measurement

The synthesized intact and fluorescent dye-labeled peptides (F-peptides, 1 mg) were dissolved in 1 mL water. A 100 μL of each sample was loaded on a black 96-well plate and subjected to fluorescence measurement. The fluorescence was scanned from 505 nm to 650 nm under 488 nm light excitation.

### 4.7. Direct peptide affinity test

Diverse concentrations of EpCAM protein (0, 1, 3, 10, 30, 100, 300, and 1,000 ng/mL) and CD81 protein (0.1, 0.3, 1, 3, 10, 30, and 100 ng/mL) were incubated overnight in each well of a black 96-well plate. The EpCAM and CD81-fixed wells were washed with 1x PBS, and then, a Superblock solution was used for surface blocking. After 2 hour-incubation and triple washing with 1x PBS, the F-peptides recognition and four 5[6]-FAM-labeled control peptides (CPs) (1 μg/mL, 100 μL) such as CP1:QWWIRNEIS, CP2:NEHRLRWIW, CP3:QFRMWEENW, and CP4:DYTWDRNYF were mixed with 0.1 % BSA and added to each well. All the probe-reacted wells were strictly washed following a 30-minute incubation in a dark room. Afterward, fluorescence at 530 nm was acquired with a spectrometer.

### 4.8. Single EV imaging

#### 4.8.1. EV isolation

Three kinds of cells (SW-620, SW620, MCF7, and MDA-MB-231) were cultured in a complete medium until they were 80%–90% confluent. Then, the cells were incubated in a medium with 2% exosome-depleted FBS (Thermo Fisher), 100 U/mL penicillin, and 100 μg/mL streptomycin for 48 h, followed by EV collection by size exclusion chromatography (SEC). the conditioned medium was collected with a 40 μm-sized cell strainer (Corning) and spun at 300 x g for 5 min to remove the cell debris. The supernatant was filtered through a 0.8 μm membrane filter (Millipore Sigma) and spun at 3,500 x g for 30 min at 4 °C by using Centricon Plus-70 Centrifugal Filter (MWCO=10 kDa, Millipore Sigma). The concentrated medium was passed through with SEC, and 4th and 5th fractions were used for EV isolation, followed by concentration using Amicon Ultra-2 Centrifugal Filter (MWCO=10 kDa, Millipore Sigma) and centrifuged at 3,500 x g for 30min at 4 °C.

#### 4.8.2. Preparation of the EV-fixed substrate

Three microliters of input samples were mixed with 2 μL of 0.1M Na_2_CO_3_ buffer, and the mixture was incubated with 0.2 μL of NHS ester-Alexa-555 (10 mg/mL in DMSO; #A37571, Thermofisher Scientific) for 1 hour at RT. Excess dyes were removed *via* washing with Zeba micro spin desalting columns (#87765, Thermofisher Scientific). After two times of washing, 3 μL of dye-labeled samples were loaded on a glass slide and allowed to settle (30 min, RT). The slide was then washed with filtered-PBS (fPBS), incubated with 10 μL of fixation buffer (4% paraformaldehyde), and washed. The revealed surface was then blocked using 15 µL of Superblock solution for 1 hour at RT.

#### 4.8.3. Bioreceptor/EpCAM protein binding on the EVs

For one-step antibody-based EpCAM detection, an anti-EpCAM antibody was incubated with NHS ester-Alexa-488 dye (10 μg/mL) for 1.5 hours and washed in fPBS. The prepared fluorescent dye-labeled antibody (F-antibody) and the F-peptide were added to the EV-fixed substrate and incubated for 1 hour. After triple washing with fPBS, fluorescence images were taken by a confocal microscope. Image analyses were then performed using ImageJ software.

## Acknowledgments

This work was supported by funds from the National Science Foundation (grant CBET- 1934284 and grant CBET-2347712). This research used resources from the ACCESS program (formerly XSEDE), supported by National Science Foundation and the North Carolina State University supercomputer Hazel. The authors also thank the San Diego Supercomputer Center (SDSC) for the computing time.

---

**Figure S1:**
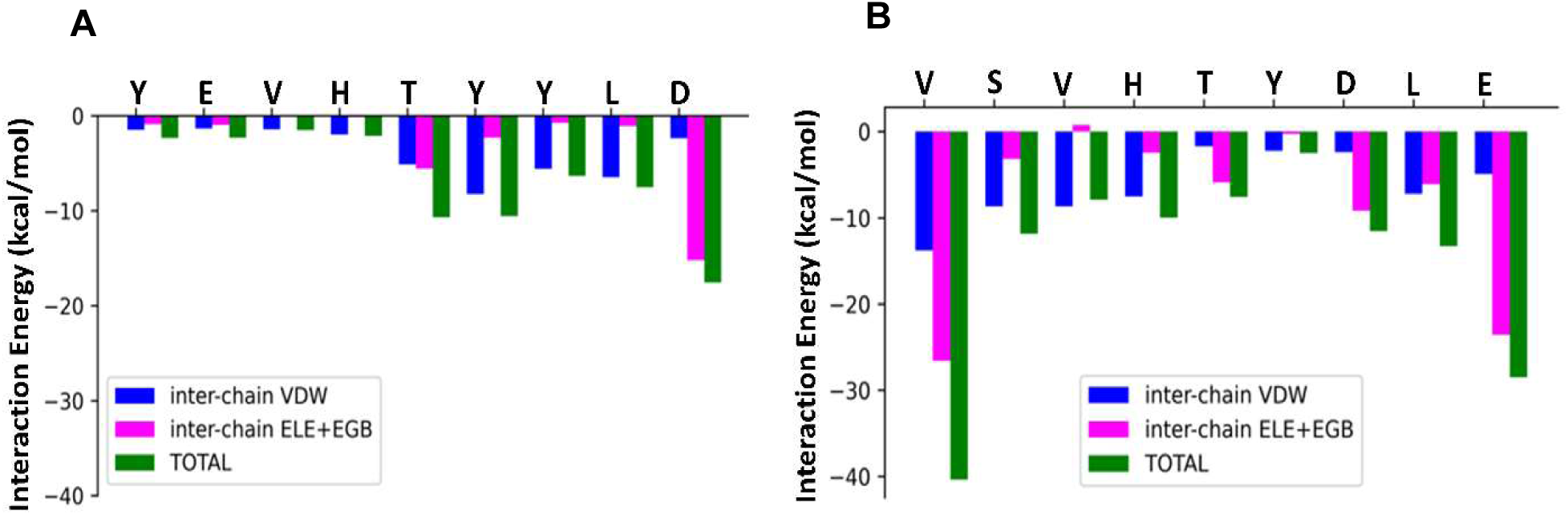
(A) The residue wise decomposition of the interaction energy plot of peptide EP- 1:EpCAM NTD and (B) the residue wise decomposition of the interaction energy plot of peptide EP-2:EpCAM P1.

**Figure S2:**
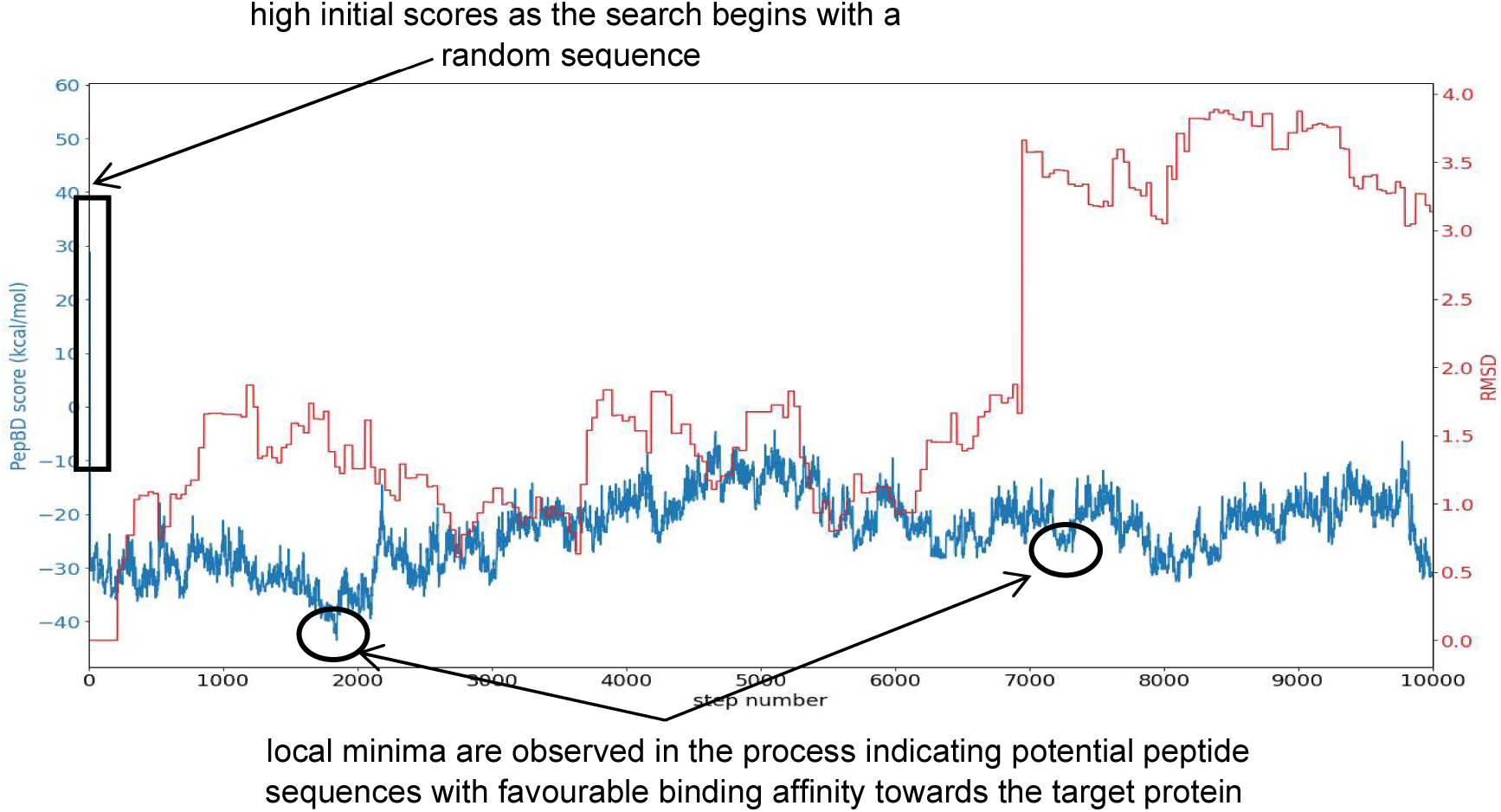
PepBD score versus the step number in blue. RMSD evolution is depicted in red. This is a typical score profile with high scores at the start of the PepBD search. The algorithm discovers new peptide sequences and generates corresponding PepBD scores, which are calculated by the score function. Low scores indicate potential peptide sequences with favorable binding free energy.

**Table S1:**
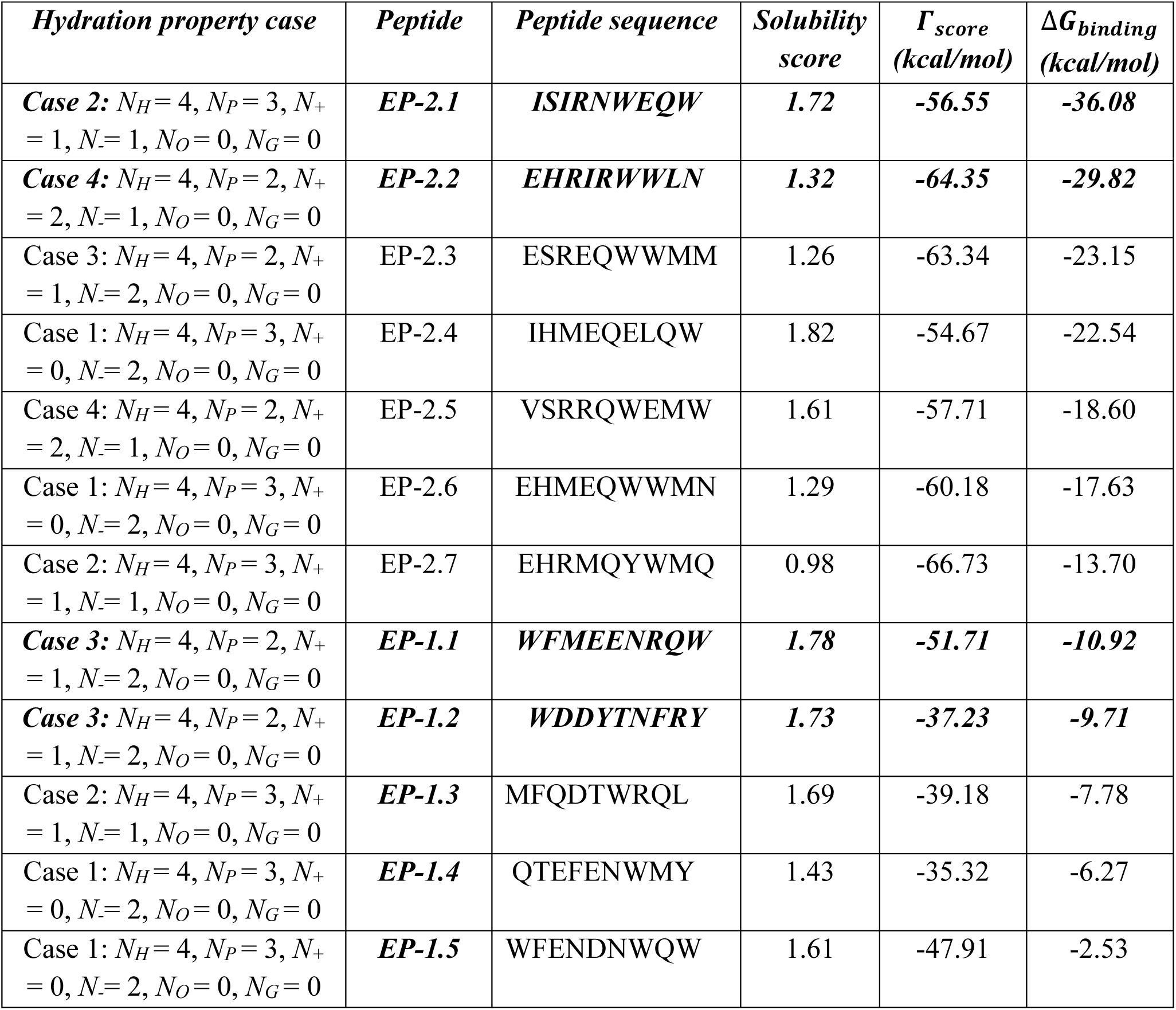
List of peptide sequences identified by PepBD screening with their corresponding 𝛤_score_ and Δ𝐺_binding_ values. The peptide sequences EP-2.1, EP-2.2 EP-1.1 and EP-1.2 (in bold and italics) are selected were selected for synthesis and experimental validation

**Table S2:**
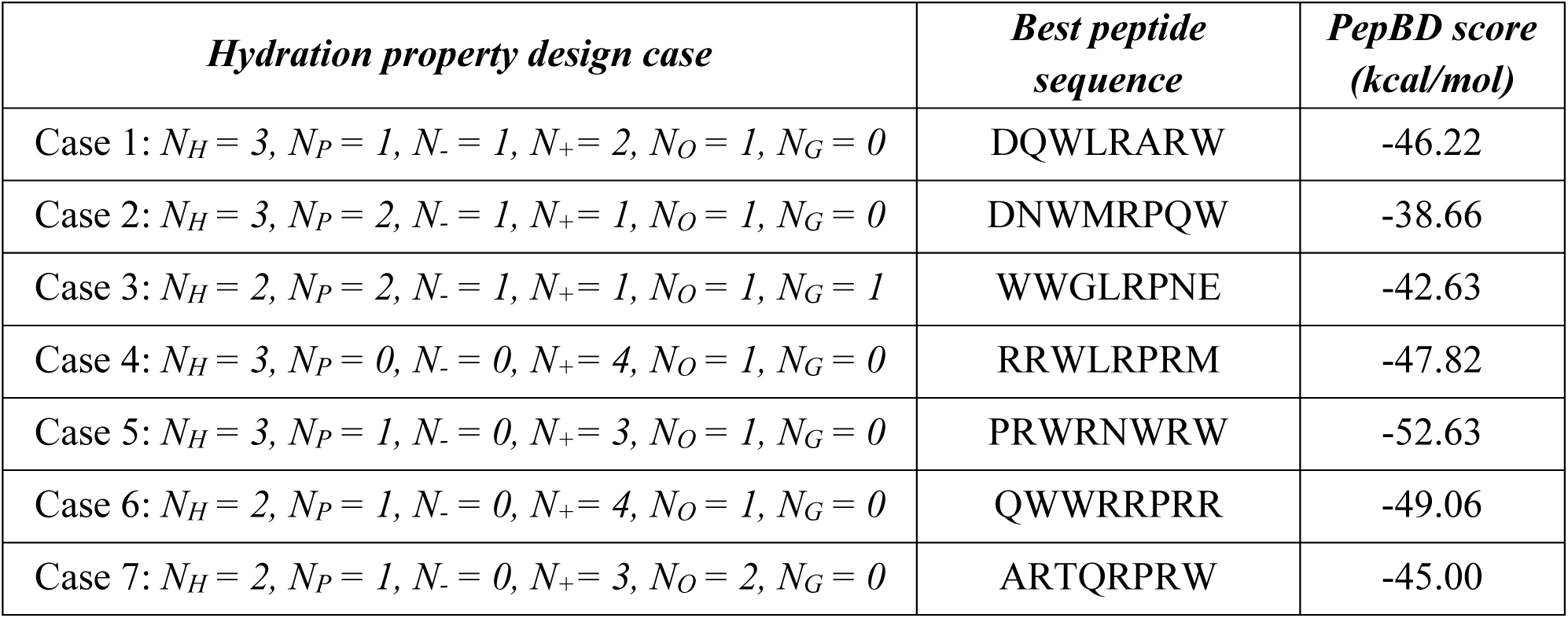
Top scoring peptide from each hydration property case along with their PepBD scores in (kcal/mol) as calculated using the scoring function 𝜞_𝒔𝒄𝒐𝒓𝒆_ for **first round** of peptide design for CD81. *N_H_, N_P_, N_-_, N_+_, N_O_, N_G_* each represent the number of hydrophobic, hydrophilic, negative, positive, other and glycine residues in the peptide sequence.

**Table S3:**
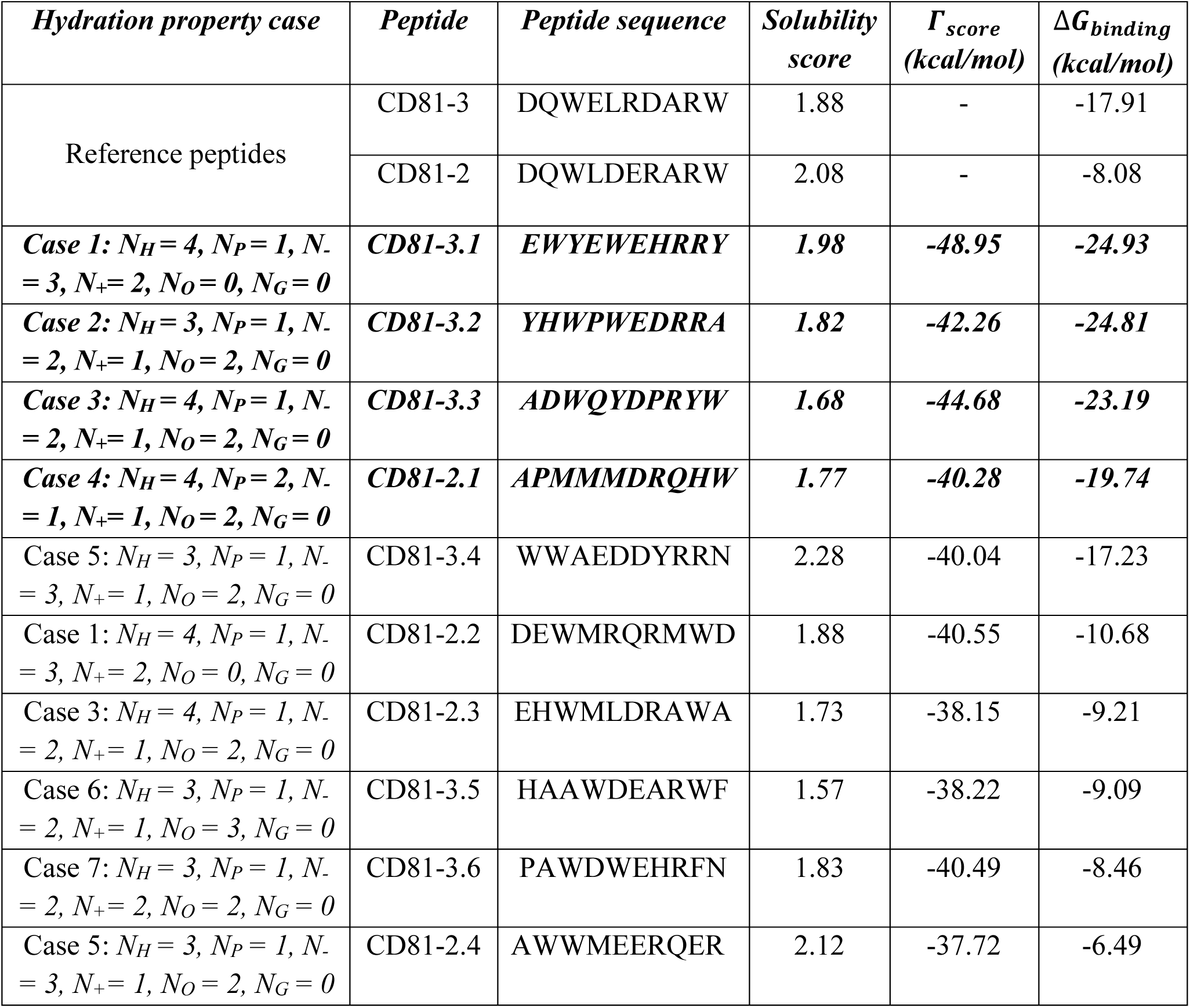
Final set of designed peptide ligands for the LEL domain of CD81. The top scoring peptide sequences are listed along with their PepBD scores, intrinsic solubility scores as calculated through the CamSol method, and best binding free energies (Δ𝐺_𝑏𝑖𝑛𝑑𝑖𝑛𝑔_ in kcal/mol) computed from running three independent explicit-solvent MD simulations for 100ns. The peptide sequences in bold and italics were selected for synthesis and experimental validation because of their low scores as compared to the other peptides.

