## Supplementary Table S1 for "Designed Peptides as Affinity Ligands for Extracellular-Vesicle-based Cancer Biomarker Detection"

**Supplementary Information (SI)**

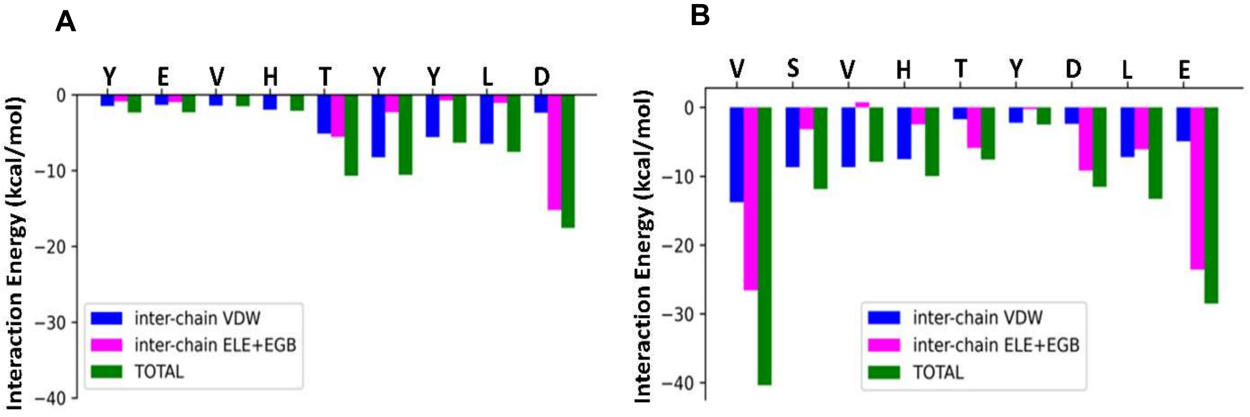


**Figure S1:** (A) The residue wise decomposition of the interaction energy plot of peptide EP-1:EpCAM NTD and (B) the residue wise decomposition of the interaction energy plot of peptide EP-2:EpCAM P1.

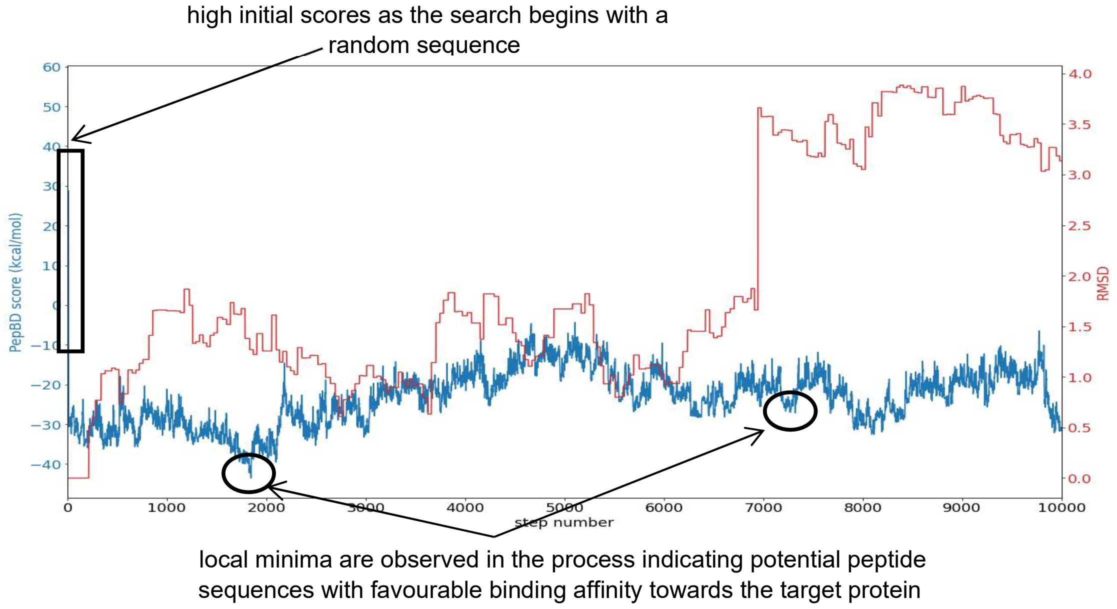


**Figure S2**: PepBD score versus the step number in blue. RMSD evolution is depicted in red. This is a typical score profile with high scores at the start of the PepBD search. The algorithm discovers new peptide sequences and generates corresponding PepBD scores, which are calculated by the score function. Low scores indicate potential peptide sequences with favorable binding free energy.

**Table S1**: List of peptide sequences identified by PepBD screening with their corresponding $\Gamma_{score}$ and $\Delta G_{binding}$ values. The peptide sequences EP-2.1, EP-2.2 EP-1.1 and EP-1.2 (in bold and italics) are selected were selected for synthesis and experimental validation

| ***Hydration property case*** | ***Peptide*** | ***Peptide sequence*** | ***Solubility score*** | $\Gamma_{score}$ ***(kcal/mol)*** | $\Delta G_{binding}$ ***(kcal/mol)*** |
|---|---|---|---|---|---|
| **Case 2:** $N_H = 4, N_P = 3, N_+ = 1, N_- = 1, N_O = 0, N_G = 0$ | **EP-2.1** | **ISIRNWEQW** | **1.72** | **-56.55** | **-36.08** |
| **Case 4:** $N_H = 4, N_P = 2, N_+ = 2, N_- = 1, N_O = 0, N_G = 0$ | **EP-2.2** | **EHRIRWWLN** | **1.32** | **-64.35** | **-29.82** |
| Case 3: $N_H = 4, N_P = 2, N_+ = 1, N_- = 2, N_O = 0, N_G = 0$ | EP-2.3 | ESREQWWMM | 1.26 | -63.34 | -23.15 |
| Case 1: $N_H = 4, N_P = 3, N_+ = 0, N_- = 2, N_O = 0, N_G = 0$ | EP-2.4 | IHMEQELQW | 1.82 | -54.67 | -22.54 |
| Case 4: $N_H = 4, N_P = 2, N_+ = 2, N_- = 1, N_O = 0, N_G = 0$ | EP-2.5 | VSRRQWEMW | 1.61 | -57.71 | -18.60 |
| Case 1: $N_H = 4, N_P = 3, N_+ = 0, N_- = 2, N_O = 0, N_G = 0$ | EP-2.6 | EHMEQWWMN | 1.29 | -60.18 | -17.63 |
| Case 2: $N_H = 4, N_P = 3, N_+ = 1, N_- = 1, N_O = 0, N_G = 0$ | EP-2.7 | EHRMQYWMQ | 0.98 | -66.73 | -13.70 |
| **Case 3:** $N_H = 4, N_P = 2, N_+ = 1, N_- = 2, N_O = 0, N_G = 0$ | **EP-1.1** | **WFMEENRQW** | **1.78** | **-51.71** | **-10.92** |
| **Case 3:** $N_H = 4, N_P = 2, N_+ = 1, N_- = 2, N_O = 0, N_G = 0$ | **EP-1.2** | **WDDYTNFRY** | **1.73** | **-37.23** | **-9.71** |
| Case 2: $N_H = 4, N_P = 3, N_+ = 1, N_- = 1, N_O = 0, N_G = 0$ | **EP-1.3** | MFQDTWRQL | 1.69 | -39.18 | -7.78 |
| Case 1: $N_H = 4, N_P = 3, N_+ = 0, N_- = 2, N_O = 0, N_G = 0$ | **EP-1.4** | QTEFENWMY | 1.43 | -35.32 | -6.27 |
| Case 1: $N_H = 4, N_P = 3, N_+ = 0, N_- = 2, N_O = 0, N_G = 0$ | **EP-1.5** | WFENDNWQW | 1.61 | -47.91 | -2.53 |

**Table S2:** Top scoring peptide from each hydration property case along with their PepBD scores in (kcal/mol) as calculated using the scoring function $\boldsymbol{\Gamma_{score}}$ for **first round** of peptide design for CD81. $N_H$, $N_P$, $N_-$, $N_+$, $N_O$, $N_G$ each represent the number of hydrophobic, hydrophilic, negative, positive, other and glycine residues in the peptide sequence.

| ***Hydration property design case*** | ***Best peptide sequence*** | ***PepBD score (kcal/mol)*** |
|---|---|---|
| Case 1: $N_H = 3, N_P = 1, N_- = 1, N_+ = 2, N_O = 1, N_G = 0$ | DQWLRARW | -46.22 |
| Case 2: $N_H = 3, N_P = 2, N_- = 1, N_+ = 1, N_O = 1, N_G = 0$ | DNWMRPQW | -38.66 |
| Case 3: $N_H = 2, N_P = 2, N_- = 1, N_+ = 1, N_O = 1, N_G = 1$ | WWGLRPNE | -42.63 |
| Case 4: $N_H = 3, N_P = 0, N_- = 0, N_+ = 4, N_O = 1, N_G = 0$ | RRWLRPRM | -47.82 |
| Case 5: $N_H = 3, N_P = 1, N_- = 0, N_+ = 3, N_O = 1, N_G = 0$ | PRWRNWRW | -52.63 |
| Case 6: $N_H = 2, N_P = 1, N_- = 0, N_+ = 4, N_O = 1, N_G = 0$ | QWWRRPRR | -49.06 |
| Case 7: $N_H = 2, N_P = 1, N_- = 0, N_+ = 3, N_O = 2, N_G = 0$ | ARTQRPRW | -45.00 |

**Table S3**: Final set of designed peptide ligands for the LEL domain of CD81. The top scoring peptide sequences are listed along with their PepBD scores, intrinsic solubility scores as calculated through the CamSol method, and best binding free energies ($\Delta G_{binding}$ in kcal/mol) computed from running three independent explicit-solvent MD simulations for 100ns. The peptide sequences in bold and italics were selected for synthesis and experimental validation because of their low scores as compared to the other peptides.

| ***Hydration property case*** | ***Peptide*** | ***Peptide sequence*** | ***Solubility score*** | $\Gamma_{score}$ ***(kcal/mol)*** | $\Delta G_{binding}$ ***(kcal/mol)*** |
|---|---|---|---|---|---|
| Reference peptides | CD81-3 | DQWELRDARW | 1.88 | - | -17.91 |
| | CD81-2 | DQWLDERARW | 2.08 | - | -8.08 |
| ***Case 1:*** $N_H = 4, N_P = 1, N_- = 3, N_+ = 2, N_O = 0, N_G = 0$ | ***CD81-3.1*** | ***EWYEWEHRRY*** | ***1.98*** | ***-48.95*** | ***-24.93*** |
| ***Case 2:*** $N_H = 3, N_P = 1, N_- = 2, N_+ = 1, N_O = 2, N_G = 0$ | ***CD81-3.2*** | ***YHWPWEDRRA*** | ***1.82*** | ***-42.26*** | ***-24.81*** |
| ***Case 3:*** $N_H = 4, N_P = 1, N_- = 2, N_+ = 1, N_O = 2, N_G = 0$ | ***CD81-3.3*** | ***ADWQYDPRYW*** | ***1.68*** | ***-44.68*** | ***-23.19*** |
| ***Case 4:*** $N_H = 4, N_P = 2, N_- = 1, N_+ = 1, N_O = 2, N_G = 0$ | ***CD81-2.1*** | ***APMMMDRQHW*** | ***1.77*** | ***-40.28*** | ***-19.74*** |
| Case 5: $N_H = 3, N_P = 1, N_- = 3, N_+ = 1, N_O = 2, N_G = 0$ | CD81-3.4 | WWAEDDYRRN | 2.28 | -40.04 | -17.23 |
| Case 1: $N_H = 4, N_P = 1, N_- = 3, N_+ = 2, N_O = 0, N_G = 0$ | CD81-2.2 | DEWMRQRMWD | 1.88 | -40.55 | -10.68 |
| Case 3: $N_H = 4, N_P = 1, N_- = 2, N_+ = 1, N_O = 2, N_G = 0$ | CD81-2.3 | EHWMLDRAWA | 1.73 | -38.15 | -9.21 |
| Case 6: $N_H = 3, N_P = 1, N_- = 2, N_+ = 1, N_O = 3, N_G = 0$ | CD81-3.5 | HAAWDEARWF | 1.57 | -38.22 | -9.09 |
| Case 7: $N_H = 3, N_P = 1, N_- = 2, N_+ = 2, N_O = 2, N_G = 0$ | CD81-3.6 | PAWDWEHRFN | 1.83 | -40.49 | -8.46 |
| Case 5: $N_H = 3, N_P = 1, N_- = 3, N_+ = 1, N_O = 2, N_G = 0$ | CD81-2.4 | AWWMEERQER | 2.12 | -37.72 | -6.49 |
